# CEBPA repression by MECOM blocks differentiation to drive aggressive leukemias

**DOI:** 10.1101/2024.12.30.630680

**Authors:** Travis J. Fleming, Mateusz Antoszewski, Sander Lambo, Michael C. Gundry, Riccardo Piussi, Lara Wahlster, Sanjana Shah, Fiona E. Reed, Kevin D. Dong, Joao A. Paulo, Steven P. Gygi, Claudia Mimoso, Seth R. Goldman, Karen Adelman, Jennifer A. Perry, Yana Pikman, Kimberly Stegmaier, Maria N. Barrachina, Kellie R. Machlus, Volker Hovestadt, Andrea Arruda, Mark D. Minden, Richard A. Voit, Vijay G. Sankaran

## Abstract

Acute myeloid leukemias (AMLs) have an overall poor prognosis with many high-risk cases co-opting stem cell gene regulatory programs, yet the mechanisms through which this occurs remain poorly understood. Increased expression of the stem cell transcription factor, MECOM, underlies one key driver mechanism in largely incurable AMLs. How MECOM results in such aggressive AML phenotypes remains unknown. To address existing experimental limitations, we engineered and applied targeted protein degradation with functional genomic readouts to demonstrate that MECOM promotes malignant stem cell-like states by directly repressing pro-differentiation gene regulatory programs. Remarkably and unexpectedly, a single node in this network, a MECOM-bound *cis*-regulatory element located 42 kb downstream of the myeloid differentiation regulator *CEBPA*, is both necessary and sufficient for maintaining MECOM-driven leukemias. Importantly, targeted activation of this regulatory element promotes differentiation of these aggressive AMLs and reduces leukemia burden *in vivo*, suggesting a broadly applicable differentiation-based approach for improving therapy.

## Introduction

Acute myeloid leukemia (AML) is an aggressive blood cancer that is curable in less than 30% of cases^1^. The inability to cure the majority of patients with AML is attributable to leukemia heterogeneity, particularly regarding the diversity of cytogenetic abnormalities, oncogenic driver mutations, and cell types of origin, all of which contribute to variable and often poor responses to standard-of-care chemotherapy^2,3^. Recently, the field has gained an appreciation for not just the identity and distribution of AML driver mutations, but also the corresponding cell state changes that alter the behavior of these cancers. Increasing evidence suggests that acquiring or persistence of hematopoietic stem cell (HSC) gene expression programs in AML confers a particularly poor prognosis and substantially increases the risk of relapse^4–7^.

These insights accompany the clinical development of AML therapies such as venetoclax^8^, menin inhibitors^9^, and preclinical candidates targeting cell death pathways, metabolic vulnerabilities, and transcriptional dependencies that seek to target leukemia stem cells^10,11^. However, many of these therapies focus on promoting death of stem cell-like populations, rather than re-establishing differentiation programs, though some such as menin inhibitors have also demonstrated activity in promoting differentiation^9^. The use of all-trans retinoic acid for treatment of *PML::RARA* fusion acute promyelocytic leukemia offers a well-established paradigm - promoting AML differentiation to enable more effective therapy^12^. However, beyond the fortuitous discovery of retinoids for acute promyelocytic leukemia, there is a limited understanding of approaches that could enable differentiation for therapeutic purposes in other subtypes of AML.

Acquisition of HSC gene expression programs in a subset of AML with particularly poor clinical prognosis^13^ is frequently driven by increased expression of *MECOM*, a transcription factor that plays a key role in normal HSC maintenance and self-renewal^14^. A number of studies have sought to elucidate the mechanisms by which MECOM drives stem cell-like, high-risk features in AML^15–19^. However, MECOM perturbation leads to an acute disruption of stem cell maintenance. As a result, molecular studies attempting to dissect MECOM’s direct role in leukemia using traditional CRISPR/Cas9 or shRNA-based loss-of-function approaches have been confounded by secondary changes due to alterations in cell state. Without a direct functional understanding of how MECOM controls stem cell regulatory networks in AML, the ability to develop mechanism-based targeted therapies has not been possible.

To address this existing limitation and investigate the role of MECOM in enabling stem cell gene expression programs to be adopted in AML, we have applied targeted protein degradation to characterize MECOM’s direct functions in a precise and temporally controlled manner. Remarkably and unexpectedly, although MECOM has a variety of targets, we uncover a previously unappreciated and simple regulatory logic underlying MECOM’s role in promoting stem cell-like states in high-risk AMLs. Repression of a single *CEBPA cis*-regulatory element by MECOM is necessary and sufficient to confer high-risk stem cell-like states in a variety of AMLs. Notably, transient activation of this *cis*-regulatory element promotes differentiation of primary leukemia cells and significantly reduces leukemia burden in *in vivo* models. These observations serve as a key proof-of-principle for the mechanistically-driven therapeutic opportunity to promote differentiation in high-risk AMLs.

## Results

### Rapid and specific protein degradation enables direct interrogation of MECOM function in AML

To directly elucidate MECOM-driven transcriptional and epigenetic programs in stem cell-like leukemia cells, we engineered three AML cell line models with a 2xHA-FKBP12^F36V^-P2A-eGFP cassette at the C-terminus of the endogenous *MECOM* locus (**Fig. 1A**). The synthetic FKBP12^F36V^ degron has been shown to enable selective and rapid degradation of tagged proteins with the addition of dTAG small molecules^20,21^. We selected the AML cell lines MUTZ-3, UCSD-AML1, and HNT-34 cells given their high *MECOM* expression level and cytogenetic status that arises due to an oncogenic translocation/inversion event that juxtaposes an enhancer of *GATA2* to drive high-level *MECOM* expression^22,23^. Consistent with *MECOM* expression being restricted to stem cell-like populations^14^, the GFP^+^ expression in these models is strongly enriched in the CD34^+^ compartment (**Fig. 1B**). Treatment of MUTZ-3 *MECOM-FKBP12*^*F36V*^ (hereafter referred to as MUTZ3-dTAG), UCSD-AML1-dTAG, and HNT-34-dTAG cells with low nanomolar (5-500 nM) concentrations of dTAG^V^-1 resulted in rapid degradation of all MECOM protein within 1 hour of treatment compared to DMSO vehicle controls (**Fig. 1C, Fig. S1A**). Moreover, multiplexed quantitative mass spectrometry demonstrated that MECOM was the only protein whose abundance was significantly altered in the proteome of MUTZ-3 cells following the addition of dTAG^V^-1 for 2 hours (fold change < −1.0, p-value < 0.001) (**Fig. 1D**). To further corroborate the specificity of this approach to rapidly ablate MECOM, we measured MECOM chromatin occupancy in CD34^+^ MUTZ-3 progenitor cells treated with dTAG^V^-1 vs.

**Figure 1:**
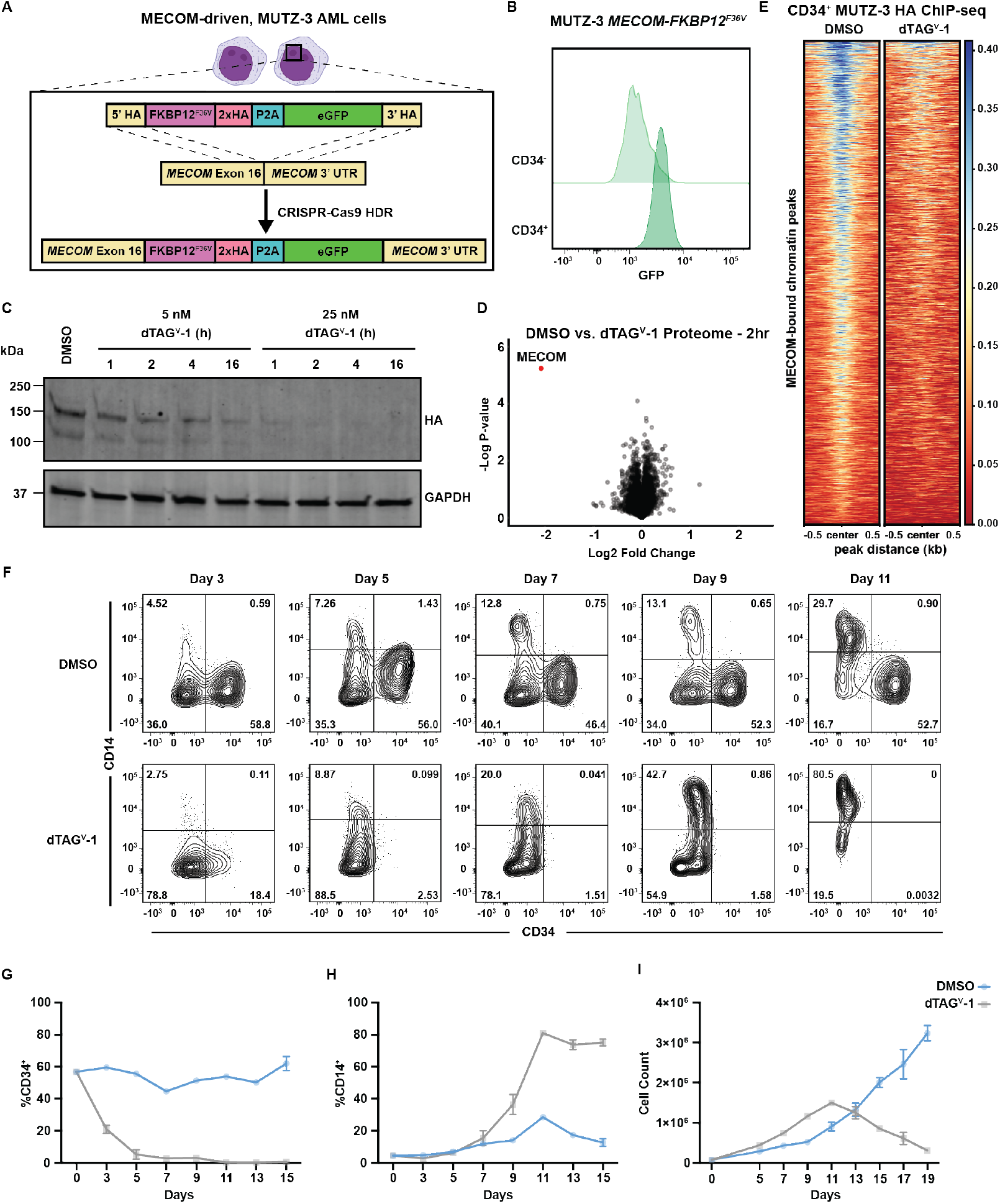
FKBP12^F36V^ degron facilitates rapid degradation of endogenous MECOM in AML cells. **(A)** Schematic illustrating the gene-editing strategy to knock-in an FKBP12^F36V^ degron, 2xHA tag, and eGFP at the C-terminus of the endogenous *MECOM* locus in human MUTZ-3 AML cells. **(B)** GFP expression assessed by flow cytometry in CD34+ vs. CD34-MUTZ-3 MECOM-FKBP12^F36V^ cells. **(C)** Time course western blot analysis of MECOM protein levels in MUTZ-3 cells following treatment with dTAG^V^-1 (5-25nM) or DMSO. **(D)** Volcano plot showing changes in protein abundance in MUTZ-3 MECOM-FKBP12^F36V^ cells treated for 2 hrs with 500nM dTAG^V^-1 vs. DMSO as assessed by mass spectrometry. n = 3 independent replicates. **(E)** MECOM ChIP-seq of MUTZ-3 MECOM-FKBP12^F36V^ cells treated with 500nM dTAG^V^-1 vs. DMSO (n=3). Each row represents a single MECOM(HA)-bound peak. Heatmap is centered on ChIP-peak summits +/−500bp. **(F)** Bivariate plot showing CD34 and CD14 expression levels in MUTZ-3 MECOM-FKBP12^F36V^ cells treated with 500nM dTAG^V^-1 vs. DMSO. **(G-H)** Percentage of CD34+ and CD14+ cells as observed in **Fig. 1F**. n = 3 independent replicates, mean and SEM are shown. **(I)** Viable cell count by trypan blue exclusion of MUTZ-3 MECOM-FKBP12^F36V^ cells treated with 500nM dTAG^V^-1 vs. DMSO. n = 3 independent replicates, mean and SEM are shown.

DMSO vehicle control and observed almost complete loss of MECOM binding across the genome (Fig. 1E). We next sought to validate the utility of these degron models to glean insights into the regulation of stem cell gene regulatory programs. Consistent with studies that genetically perturb *MECOM*^14^, MUTZ-3-dTAG cells treated with dTAG^V^-1 exhibited nearly complete loss of CD34 expression followed by acquisition of CD14 expression consistent with monocytic differentiation (**Fig. 1F-H**). Although dTAG^V^-1-treated MUTZ-3 cells initially proliferate more, they eventually all die in culture presumably due to loss of stem cell/progenitor populations and the short persistence of terminally differentiated cells^14,24^ (**Fig. 1I**). This robust myeloid differentiation phenotype was conserved in UCSD-AML1-dTAG cells, where dTAG^V^-1 treatment resulted in loss of CD34 expression and morphological signs of differentiation (**Fig. S1 B-D**). We did not observe signs of morphologic or immunophenotypic differentiation in HNT-34-dTAG cells following MECOM degradation, however, the cells rapidly underwent apoptosis in culture (**Fig. S1 E-F**). This is in agreement with a previous report describing a strong MECOM dependency in HNT-34 cells^19^. To further profile the impact of synchronous loss of MECOM, we utilized a fluorescent EdU-labeling assay to analyze cell cycle differences induced upon MECOM loss. dTAG^V^-1 treatment conferred a significant increase in actively dividing cells in S and G2 phases and a significant decrease in cells in G0/G1 phase (**Fig. S1 G-H**). This is consistent with the differentiation phenotypes observed, as loss of quiescence accompanies hematopoietic differentiation. Together, these experiments demonstrate the utility of the dTAG system to rapidly and specifically degrade MECOM in cellular models of leukemia. Importantly, these models enable sensitive molecular profiling following MECOM ablation and prior to cell state changes, which is crucial for elucidating its direct role in enabling stem cell phenotypes in high-risk AMLs.

### MECOM directly represses myeloid differentiation programs in AML

Having established and validated several MECOM degron models, we next performed multiomic profiling following MECOM degradation to elucidate regions of accessible chromatin and genes directly regulated by this factor. We reasoned that by restricting our profiling to stem cell-like, MECOM-expressing cells we would enhance our ability to detect direct transcriptional and epigenetic alterations. Following CD34^+^ enrichment (**Fig. 2A**), MUTZ-3 cells were treated with 500 nM dTAG^V^-1 and analyzed for changes in nascent transcription via Precision Run-On Sequencing (PRO-seq), bulk transcription (bulk RNA-seq), and chromatin accessibility (ATAC-seq). One hour after MECOM degradation we detected both increases (468 genes, p < 0.01, Log2FoldChange (L2FC) > 0.5) and decreases (600 genes, p < 0.01, L2FC < 0.5) in nascent gene expression (**Fig. 2B, Table S3**). However, by four hours (PRO-seq) (**Fig. 2C**) and six hours (bulk RNA-seq) (**Fig. 2D, Table S3-4**) after MECOM degradation far more genes showed increased expression (4hr: 153 increased vs. 64 decreased genes, 6hr: 47 increased vs. 8 decreased genes, respectively), suggestive of a direct repressive function for MECOM in this context. Moreover, a recent study from our group elucidated an HSC gene signature that is downregulated upon MECOM perturbation in primary human HSCs (MECOM down genes)^14^. Notably, far more “MECOM down” genes show significantly reduced expression at 24 hours vs. 6 hours after MECOM degradation (58 genes vs. 1 gene, respectively, L2FC < −0.5, p < 0.001) (**Fig. S2A-D**), highlighting how the loss of stem cell maintenance gene programs in AML is likely to be secondary to the activation of myeloid differentiation programs observed upon acute loss of MECOM.

**Figure 2:**
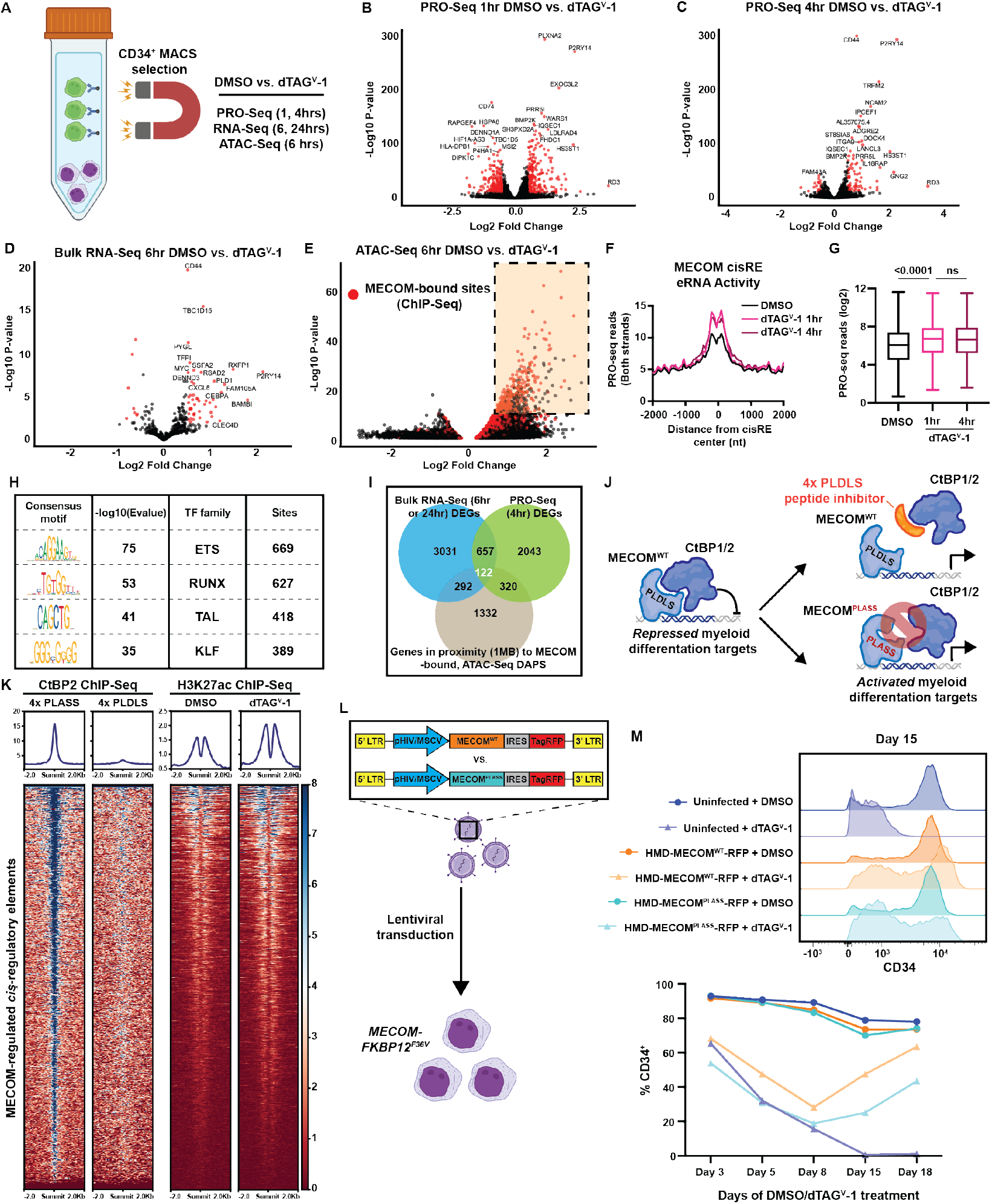
Multiomic profiling of MECOM-depleted cells reveals a predominantly repressive role at target sites. **(A)** Schematic representation of experimental protocol for multiomic characterization of dTAG^V^-1 treated MUTZ-3 MECOM-FKBP12^F36V^ cells. The CD34+, GFP+ MECOM-expressing population was pre-enriched via magnetic-activated cell sorting (MACS) prior to treatment with 500nM dTAG^V^-1 or DMSO. Cells were then harvested and processed for bulk RNA-seq, ATAC-seq, and Precision run-on sequencing (PRO-seq) to profile transcriptional and epigenetic changes. **(B-C)** Volcano plots representing changes in nascent gene expression assessed via PRO-seq in MUTZ-3 MECOM-FKBP12^F36V^ cells treated with dTAG^V^-1 vs. DMSO for 1 and 4 hours. n = 3 independent replicates. **(Table S4). (D)** Volcano plot representing changes in gene expression assessed via bulk RNA-seq in MUTZ-3 MECOM-FKBP12^F36V^ cells treated with dTAG^V^-1 vs. DMSO for 6 hours. n = 3 independent replicates **(Table S3). (E)** Volcano plot representing changes in chromatin accessibility as assessed by ATAC-seq in MUTZ-3 MECOM-FKBP12^F36V^ cells treated with dTAG^V^-1 vs. DMSO for 6 hours. n = 3 independent replicates. Red data points represent chromatin peaks that are also bound by MECOM as assessed by MECOM-HA ChIP-seq. There are 837 of these sites that are schematically highlighted in the top right corner of the plot **(Table S5). (F-G)** Assessment of enhancer RNA (eRNA) transcription levels at 837 MECOM-bound differentially accessible peaks measured from PRO-seq data. **(F)** Average PRO-seq read density across all MECOM-regulated cisREs with +/−2000bp on each side of the peak summit in dTAG^V^-1 vs. DMSO treated samples. **(G)** Box plot showing average PRO-seq read density in aggregate for each MECOM-regulated cisRE +/−500bp on each side of the peak summit in dTAG^V^-1 vs. DMSO treated samples. Two-sided Student t test was used for comparisons. n = 3 independent replicates, ns, not significant. **(H)** Unbiased motif enrichment analysis of ATAC-seq differentially accessible peaks between dTAG^V^-1 and DMSO treated samples. **(I)** Venn diagram comparing gene expression and chromatin accessibility changes across sequencing modalities. Bulk RNA-seq differentially expressed genes (DEGs) from 6hr and 24hr dTAG^V^-1 treatment, PRO-seq DEGS from 4hr dTAG^V^-1 treatment, and genes in proximity (within 1 MB) to at least one MECOM-bound, differentially accessible ATAC-seq peak were overlapped to yield a consensus MECOM gene network consisting of 122 genes. Cutoffs for bulk RNA-seq and PRO-seq were p < 0.05 **(Table S6-7)**. Peak-to-gene proximity was determined using the Genomic Regions Enrichment of Annotations Tool (GREAT)^29^. DAP, differentially accessible peak. **(J)** Schematic depiction of MECOM’s interaction with transcriptional co-repressor CtBP2 via MECOM’s PLDLS motif. This protein-protein interaction can be inhibited by a genetically-encoded 4x PLDLS peptide inhibitor^30^ (top) or if MECOM’s PLDLS motif is mutated to PLASS (bottom). **(K)** H3K27ac and CtBP2 ChIP-seq analysis. (Left) Heatmap displays CtBP2 ChIP-seq signal at MECOM-regulated cisREs in MUTZ-3 cells expressing a 4x-PLDLS peptide inhibitor of the MECOM-CtBP2 interaction compared to cells expressing 4x-PLASS control^30^. (Right) Heatmap showing H3K27ac ChIP-seq signal at MECOM-regulated cisREs in MUTZ-3 MECOM-FKBP12^F36V^ cells treated with 500nM dTAG^V^-1 or DMSO for 6 hours. **(L-M)** Experimental overview for lentiviral MECOM add-back rescue experiment. **(L)** MUTZ-3 MECOM-FKBP12^F36V^ cells were transduced with lentiviruses constitutively expressing either WT MECOM (EVI1 isoform) or MECOM PLDLS>PLASS along with a TagRFP transduction reporter at high MOI. **(M)** CD34 expression assessed by flow cytometry as a function of treatment duration (500nM DMSO vs. dTAG^V^-1) (bottom). Histogram of CD34 expression at day 15 (top). Samples were transduced 48 hours prior to treatment. n = 3 independent replicates, mean and SEM are shown but many are hidden due to low variation between replicates.

Analysis of genome-wide chromatin accessibility six hours after MECOM degradation showed a significant skew regions with towards increased accessibility, further corroborating this primarily repressive role for MECOM (**Fig. 2E, Table S5**). Specifically, accessible chromatin regions (p < 0.0001) showed strong increases in accessibility (3,071/3,462 peaks (88.7%) with L2FC > 0.5, p < 0.0001) compared to a small number of peaks with decreased accessibility (155/3,462 peaks (4.48%) with LFC < −0.5, p < 0.0001). Moreover, when overlapping these differentially accessible peaks with MECOM ChIP-seq peaks (highlighted in red), we observe a striking enrichment of overlapped peaks that show increases in accessibility (1,602/3,071 overlapping peaks; 52.2% vs. 14.4% (44,425/308,630) of ChIP-seq peaks overlapping any accessible chromatin region). We further restricted analysis to a subset of these differentially accessible sites (837) with strong MECOM chromatin occupancy (ChIP-seq P-score (−log10 p-value *10) > 50) and henceforth refer to this network of MECOM-bound sites as the direct MECOM *cis*-regulatory element (cisRE) network (**Table S6**). Across this cisRE network we detected an increase in enhancer RNA (eRNA) transcription, which serves as a proxy for enhancer activity^25,26^, following MECOM degradation (**Fig. 2F-G**). Moreover, transcription factor motif enrichment analysis of this MECOM cisRE network revealed strong enrichment of ETS motifs (**Fig. 2H**) consistent with prior reports of the binding specificity of MECOM’s C-terminal zinc finger domain^27,28^. To define a consensus, directly regulated MECOM gene signature, we integrated results from these multi-omic readouts. To do so, we first employed the Genome Regions Enrichment of Annotations Tool (GREAT)^29^ to link our MECOM cisRE network to genes by proximity. This analysis nominated 1,332 genes that were within 1 MB of at least one cisRE. We then took the union of this gene set, the differentially expressed genes from bulk RNA-seq (6 or 24hr, p < 0.05), and those obtained from PRO-seq (4hr, p < 0.05) to define a consensus network of 122 genes that might be under direct regulation of MECOM (**Fig. 2I, Table S7**). To validate that these MECOM-regulated cisRE and gene networks were conserved across multiple AML models we performed Gene Set Enrichment Analyses (GSEA) of these MECOM-regulated cisRE and gene networks in UCSD-AML1-dTAG and HNT-34-dTAG cells and showed a strong enrichment of both networks in cells treated with dTAG^V^-1 vs. DMSO via bulk RNA-seq and ATAC-seq (**Fig. S2E-L**). Collectively, these results indicate that MECOM is able to promote stem cell-like phenotypes in AML by repressing a highly conserved myeloid differentiation program.

Given these findings, we hypothesized that MECOM’s repressive role might be enabled by its interaction with transcriptional co-repressors. Indeed, MECOM has previously been shown to bind the C-terminal binding proteins 1 and 2 (CtBP1/2) through a PLDLS motif^30–32^ and this interaction can be blocked through addition of a peptide inhibitor^30^ (**Fig. 2J**). We analyzed CtBP2 ChIP-seq data^30^ from MUTZ-3 cells treated with this peptide inhibitor and observed loss of CtBP2 binding in our MECOM cisRE network consistent with a model whereby MECOM recruits CtBP2 to repress cisREs (**Fig. 2K**). Consistent with loss of CtBP2 occupancy, MECOM degradation also confers a significant increase in H3K27 acetylation across the MECOM cisRE network (**Fig. 2K**). To further validate the significance of this interaction, we performed a lentiviral rescue experiment with either exogenously expressed MECOM^WT^ or MECOM^PLDLS>PLASS^ (an isoform unable to interact with CtBP1/2) to rescue dTAG^V^-1-induced differentiation (**Fig. 2L**). In line with the importance of this interaction for the repressive function of MECOM, MECOM^WT^ rescued the loss of CD34^+^ progenitor cells induced by dTAG^V^-1 treatment of MUTZ-3 cells to a greater extent than MECOM^PLDLS>PLASS^ (**Fig. 2M**).

### MECOM gene regulatory networks are highly conserved in primary AMLs

Having identified a highly conserved network of genes repressed by MECOM that are critical for myeloid differentiation in AML cell lines, we sought to examine how conserved these programs might be in primary leukemias. We leveraged bulk transcriptome, single-cell RNA sequencing (scRNA-seq), and single-cell ATAC sequencing (scATAC-seq) from pediatric patients enrolled in the AAML1031 clinical trial^33,34^ to investigate whether gene expression and chromatin accessibility changes observed after MECOM depletion were present in patients with high *MECOM* expression, as compared to those without *MECOM* expression. Within this dataset out of 701 AML samples, 67 samples had high expression of *MECOM* (log_2_ expression > 5). The majority of leukemias expressing *MECOM* were driven by MLL rearrangements (MLLr) (59.7%), with other notable rearrangements being NUP98 fusions and MECOM fusions, confirming previously described subgroup specific patterns of MECOM expression^35^ (**Fig. S3A**). We focused on MLLr AMLs, given that this comprised the largest cohort (**Fig. S3B**). Overall survival and event-free survival was significantly worse in patients with AML with detectable/high *MECOM* expression (**Fig. S3C**) underscoring the importance of MECOM’s gene regulatory activity in relation to patient survival, as has been reported previously^36,37^. Within the cohort, *MECOM* expression also correlated with significantly higher levels of HSC-associated genes CD34 and SPINK2 (**Fig. S3D**). Similar to previous reports^38^, *MECOM* expression correlated with a scRNA-seq-derived non-malignant HSC signature^34^, further supporting the role of MECOM in enabling stem cell-like gene expression programs in AML (**Fig. S3E-F**). Given the limitations of employing bulk data, due to contaminating non-malignant cells and overall leukemia heterogeneity, we investigated differential gene expression in MLLr AMLs using single cell genomic data. Out of 11 samples sequenced in the AAML1031 cohort, five expressed MECOM and six did not express MECOM (**Fig. S3G**). To investigate whether *MECOM* expression within a leukemia correlates with the differentiation phenotype, we examined signatures from non-malignant HSCs and monocytes within each cell as well as our 122 gene signature that we had defined to be directly regulated by MECOM (**Fig. 3A-B**). Indeed, HSC signatures were significantly more abundant in AMLs expressing MECOM, while monocyte signatures were significantly more abundant in leukemias lacking MECOM. Notably, the gene signature we defined as being repressed by MECOM was lower in *MECOM* expressing AMLs (**Fig. 3C**). Furthermore, when performing differential expression analysis between leukemias expressing MECOM or not, we observed that individual HSC-associated genes are upregulated in MECOM expressing leukemias, while monocyte associated genes and MECOM network genes (none of which are monocyte associated genes) are downregulated in leukemias expressing MECOM (**Fig. 3D**). These findings confirm that the phenotypic patterns and gene expression changes that we empirically observed in our cell line models upon acute depletion of MECOM (**Fig. 2**) were recapitulated in primary AMLs.

**Figure 3:**
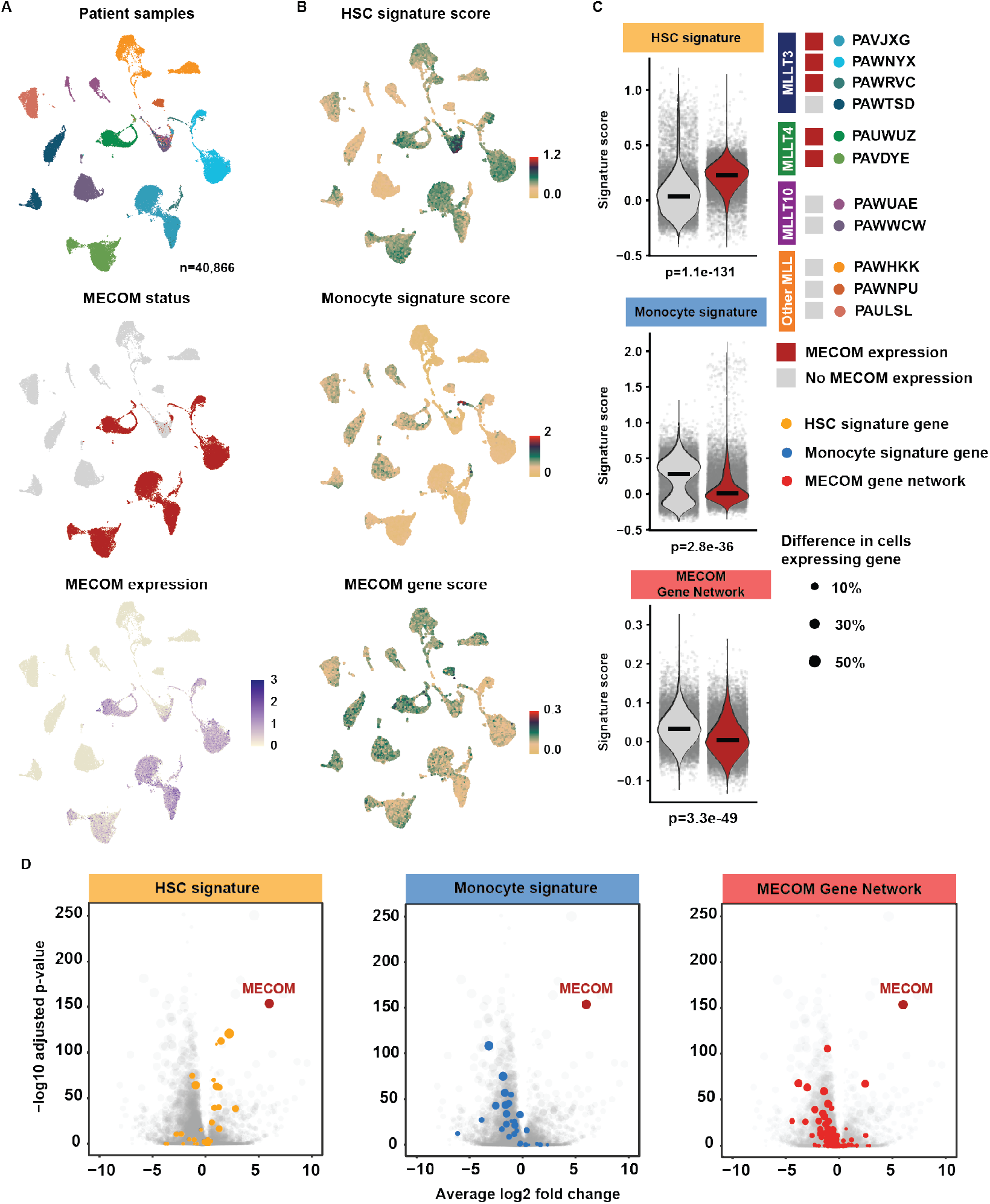
Direct MECOM gene network is repressed in primary leukemia cells. **(A)** UMAP of 40,866 cells derived from 11 patients with leukemias driven by MLL rear-rangements sequenced using single cell RNA sequencing. UMAPs were colored from top to bottom by patient, whether MECOM is expressed, and MECOM expression counts per cell. **(B)** Same UMAPS as **(A)** colored by expression signatures of normal HSCs and normal monocytes derived from Lambo et al 2023^34^ and MECOM-regulated genes identified to be activated after depletion of MECOM **(Figure 2)**. **(C)** Quantification of the three signatures from **(B)** compared between MECOM positive leukemias (n=5) and MECOM negative leukemias (n=6). Significance was calculated using two sided Wilcoxon signed-rank tests corrected for multiple testing using BH. **(D)** Differential expression of all analyzed genes (n=28,113) between leukemias expressing MECOM and leukemias that did not express MECOM. Differential expression was performed using MAST using 10 iterations of 1000 randomly selected MECOM positive cells and 1000 randomly selected MECOM negative cells to prevent uninformative p-values. BH corrected p-values and log fold changes shown are the average of 10 iterations.

### Conservation of MECOM *cis*-regulatory element network in primary AMLs

We next investigated whether the chromatin alterations observed *in vitro* after acute depletion of MECOM correlate with changes observed due to high MECOM expression in primary AMLs. Given the heterogeneity of chromatin alterations and cell states between different AMLs, we analyzed scATAC-seq of matched remission samples (n=20 patients) from the AAML1031 cohort, which expressed MECOM and had identifiable trajectories of myeloid, erythroid, and lymphoid differentiation (**Fig. 4A**). To quantify differentiation within the scATAC-seq data, we utilized peak sets that are specifically accessible in myeloid, lymphoid, and erythroid trajectories and calculated the relative peak accessibility in comparison to other trajectories. This analysis yielded differentiation scores from HSC-like cells to monocytic, mature B-cell, and late erythroid cells (**Fig. 4B**). We then calculated, for each cell, the total number of peaks identified to be repressed by MECOM (MECOM cisRE score), which we identified as being particularly abundant in cells in the HSC-like to monocyte-like axis (**Fig. 4C**). Having calculated MECOM cisRE scores and differentiation scores for each cell, we correlated the scores across all cells and found that accessible chromatin peaks identified to be bound by MECOM were correlated with myeloid differentiation scores, while they were anticorrelated with erythroid differentiation scores (**Fig. 4D**). This confirms that MECOM-regulated peaks become increasingly accessible during myeloid differentiation. To investigate the peaks in more detail we inferred a trajectory of differentiation towards monocyte-like populations from cells expressing MECOM using Monocle^39^ (**Fig. 4E, F**). We then correlated gene expression of putative MECOM target genes in linked scRNA-seq from 20 matched scRNA-seq samples and accessibility of MECOM *cis*-regulatory regions from scATAC-seq (**Fig. 4G**). We observed substantial decreases in chromatin accessibility and gene expression across the myeloid differentiation trajectory. These analyses in primary leukemia and matched remission samples corroborate MECOM’s role in directly repressing a myeloid differentiation cisRE network.

**Figure 4:**
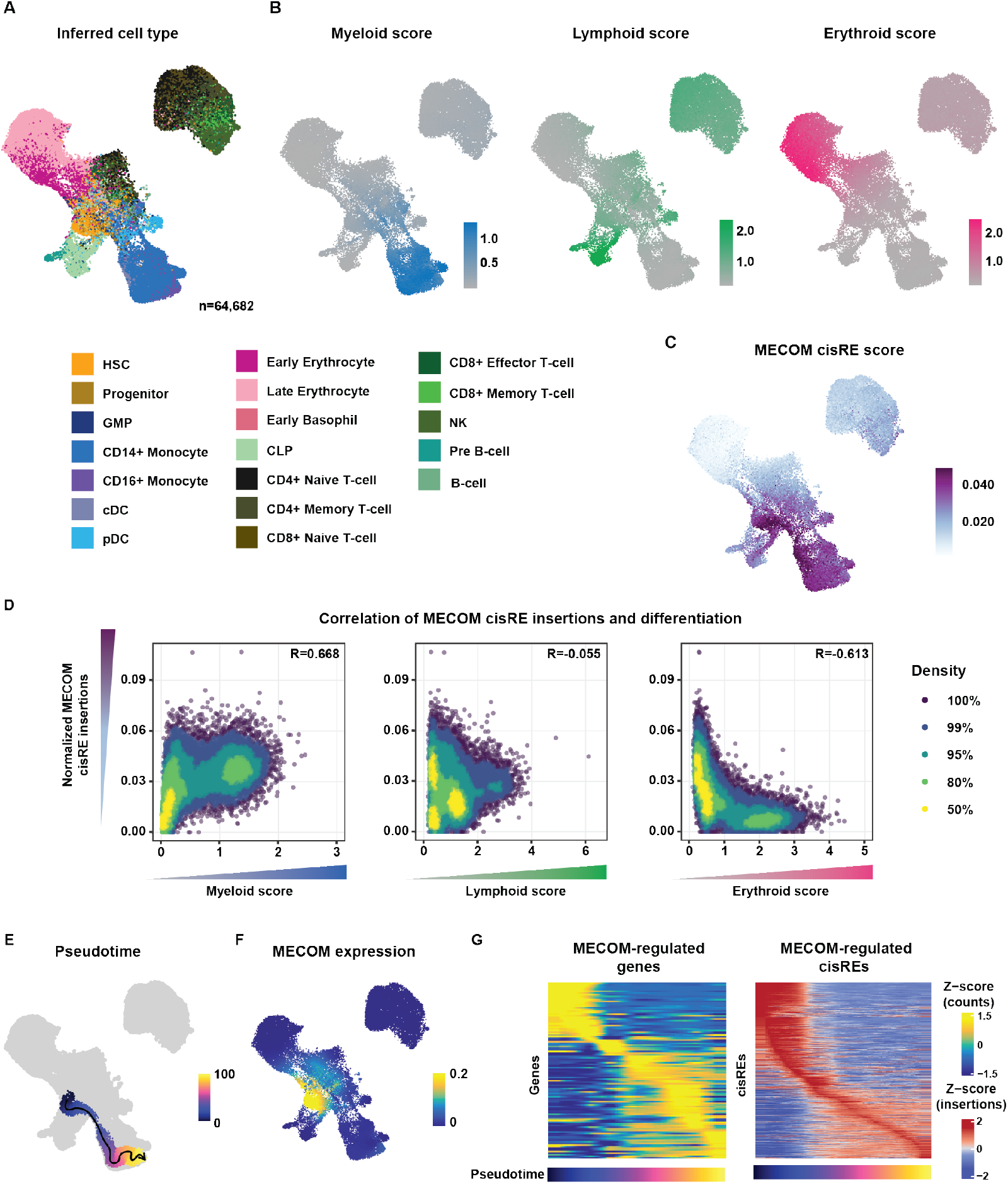
Direct MECOM chromatin network is repressed in primary leukemia cells. **(A)** UMAP of 64,682 cells generated using scATAC-seq from remissions of pediatric AML patients (n=20). Cells were colored by predicted cell type derived using label transfer of matching scRNA-seq data. Labels were derived from Lambo et al 2023^34^. **(B)** UMAP showing the same cohort as **Fig. 4A** colored by lineage scores. Lineage scores were calculated as the total insertions in 5000 accessible sites in each lineage, normalized to accessible sites in the other two lineages (accessible sites derived from Lambo et al 2023^34^). **(C)** Cells colored by chromatin accessibility at MECOM-bound loci that were identified to increase in accessibility after depletion of MECOM. Scores were calculated by ATAC-seq reads at MECOM-bound loci divided by ATAC-seq reads in the TSS and corrected for Tn5 bias. Scores were scaled to the 99th quantile to reduce the effect of outliers. **(D)** Spearman correlation between lineage scores and MECOM cisRE scores. Each dot represents one cell, cells were colored by density. **(E)** UMAP showing a trajectory inferred using Monocle from inferred HSCs to inferred Monocytes. **(F)** UMAP showing the scaled expression of MECOM in counts from linked scRNA data across cells from remissions. **(G)** Heatmaps showing the scaled expression of MECOM-regulated genes (n=122) and cisREs (n=837) along the monocyte trajectory (pseudotime). Each column represents an aggregated minibulk from cells across the inferred pseudotime (100 bins total). Normalized gene expression scores are derived from linked scRNA samples, ATAC-seq signal was normalized by TSS insertions and Tn5 bias. Both gene expression and ATAC-seq signal were scaled across all cells in the pseudotime.

In addition to these correlative analyses, we sought to empirically validate the conservation of these MECOM-driven gene and cisRE networks in an MLLr cell line model. To do so, we examined *MECOM* expression in MLLr AMLs in the Cancer Dependency Map^40^, and found high expression in OCI-AML4, an *MLL::ENL* fusion cell model^41^ (**Fig. S4A**). Therefore, we created an additional CRISPR engineered, biallelically-tagged MECOM-dTAG isogenic model in OCI-AML4 cells (**Fig. S4B**) and confirmed that dTAG^V^-1 treatment induced rapid MECOM degradation (**Fig. S4C**). We then leveraged this model to perform additional bulk RNA-seq and ATAC-seq of dTAG^V^-1-treated OCI-AML4 cells. Differential expression and gene set enrichment analyses confirmed that our repressive MECOM gene and cisRE networks are conserved in this cytogenetically distinct model of MECOM-expressing, high-risk AML (**Fig. S4D-I**). Overall, our empiric and correlative analyses of MECOM gene regulatory functions illustrate how MECOM acts as a gatekeeper of a highly conserved myeloid differentiation program across diverse high-risk AMLs.

### Functional CRISPRi/a screens identify a *CEBPA* cisRE as a key MECOM-controlled *cis*-regulatory element to block differentiation

Having defined and validated a conserved myeloid differentiation gene network repressed by MECOM, we next sought to pinpoint critical nodes in this network. Functional genomic screens were employed to identify MECOM-controlled cisREs that are essential in facilitating MECOM’s ability to block differentiation in stem cell-like leukemia cells. We designed a lentiviral sgRNA library comprised of 2,741 sgRNAs to target MECOM-repressed cisREs (**Methods**) and employed it in both a CRISPR inhibition^42^ (CRISPRi) rescue screen and a CRISPR activation^43^ (CRISPRa) differentiation screen in MUTZ-3 AML progenitor cells (**Fig. 5A**). First, in the CRISPRi screen we treated dCas9-KRAB-expressing cells with dTAG^V^-1 while simultaneously repressing individual MECOM-regulated cisREs with KRAB to investigate whether the repression of any single cisRE was sufficient to maintain leukemia cells in a CD34^+^ stem cell-like state in the absence of endogenous MECOM. After two weeks of culture, we sequenced integrated sgRNAs in the CD34^+^ phenotypically rescued population **(Fig. S5A)** and identified a strong enrichment of positive control sgRNAs targeting the transcription start sites (TSS) of *VHL, ELOB*, and *ELOC* (**Fig. 5B**). As anticipated, knockdown of VHL or the ELOB-ELOC subcomplex^44^ renders dTAG^V^-1, a VHL-targeting PROTAC, inactive. Thus, cells expressing these sgRNAs are resistant to dTAG^V^-1-mediated degradation and retain a stem cell-like phenotype, characterized by sustained CD34 expression. We also identified a significant enrichment of sgRNAs targeting both a cisRE 20 kb upstream from the *RUNX1* TSS (*RUNX1* −20 kb) and a cisRE 42 kb downstream from *CEBPA* TSS (*CEBPA* +42 kb) (**Fig. 5B, Table S8**). In an orthogonal interrogation of cisRE function, we next investigated whether activation of any single MECOM-repressed element, in the absence of MECOM perturbation, is sufficient to induce differentiation of stem cell-like leukemia cells. To accomplish this, we engineered MUTZ-3 cells to constitutively express the targetable transcriptional activator dCas9-VPR and transduced them with the same MECOM cisRE-targeting sgRNA library. Rather than sorting for CD34^+^ cells, in this screen we flow cytometrically sorted and sequenced sgRNAs from CD34^−^ cells undergoing myeloid differentiation (**Fig. 5A**). Strikingly, the only significant hit from this CRISPRa pro-differentiation screen was the same *CEBPA* +42 kb cisRE **(Fig. 5C, Table S8)**. Given that this *CEBPA*-linked cisRE was the strongest hit from both screens and has been previously implicated in myeloid differentiation^45^, we selected this target for further validation. For single sgRNA validation studies we utilized our top 3 performing *CEBPA* cisRE-targeting sgRNAs (**Fig. 5D**) and compared them to a non-targeting (sgNT) control sgRNA. In line with our pooled screening results, KRAB-mediated repression of the *CEBPA* +42 kb cisRE resulted in decreased *CEBPA* expression and maintained leukemia cells in a CD34^+^ stem cell-like state following MECOM degradation **(Fig. 5E-F, Fig. S5B)**. Furthermore, activation of this cisRE alone was sufficient to increase *CEBPA* expression and, notably, drive the differentiation of AML cells without disrupting endogenous MECOM function (**Fig. 5G-H, Fig. S5C**).

**Figure 5:**
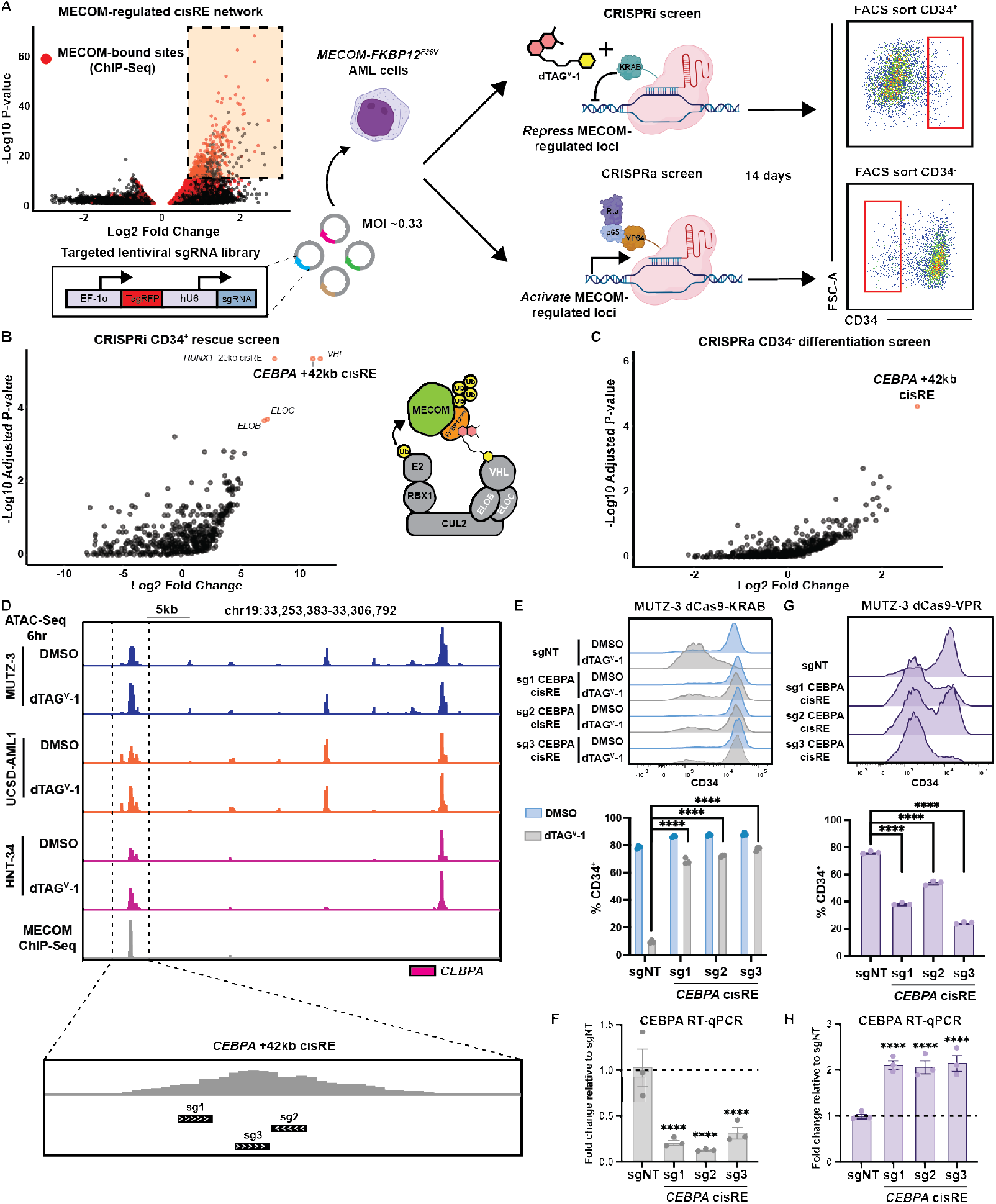
Functional CRISPR screening identifies CEBPA cisRE as a key regulator of myeloid differentiation in high-risk leukemia. **(A)** Schematic overview of the CRISPR screens utilized to functionally interrogate MECOM-regulated *cis*-regulatory elements. An sgRNA oligo library was designed against MECOM-regulated elements (up to 5 sgRNAs per element, depending on availability of high quality sgRNAs-targeting sites) and packaged into a lentiviral vector. Two different populations of MUTZ-3 MECOM-FKBP12^F36V^ cells were then transduced with this sgRNA library virus at an MOI of ~0.33; one population expressing dCas9-KRAB (CRISPRi screen) and another expressing dCas9-VPR (CRISPRa screen). Cells in the CRISPRi screen were treated with 500nM dTAG^V^-1 for the duration of the screen. After 14 days of in vitro culture, cells from the CRISPRi screen and CRISPRa screen were sorted for phenotypically rescued CD34+ cells (up-assay) and differentiated CD34-cells (down-assay), respectively. Genomically integrated sgRNAs were sequenced to assess relative sgRNA abundance. Both screens were performed with n = 3 independent replicates. **(B-C)** Volcano plots depicting sgRNA enrichment/depletion from sorted populations compared to plasmid library DNA (pDNA) **(Table S8)**. The sgRNA library included sgRNAs targeting the transcription start sites (TSS) of *VHL, ELOB*, and *ELOC* (5 sgRNAs per gene) which form the E3 ubiquitin-ligase complex recruited by dTAG^V^-1. **(D)** Genome browser tracks at the *CEBPA* locus encompassing the +42kb cisRE. ATAC-seq tracks from MECOM-FKBP12^F36V^ cell line models and MECOM ChIP-seq demonstrate increased chromatin accessibility upon dTAG^V^-1 treatment. Three top-scoring *CEBPA* cisRE-targeting sgRNAs were selected for single sgRNA validation experiments. **(E)** MUTZ-3 dCas9-KRAB cells were infected with sgRNA-expressing lentiviruses targeting either the CEBPA cisRE or a non-targeting (NT) sequence. 48 hours after transduction, cells were treated with 500nM dTAG^V^-1 vs. DMSO. (Top) Histogram shows CD34 expression at day 9. (Bottom) Percentage of CD34+ cells at day 9. n = 3 independent replicates, mean and SEM are shown. **(F)** RT-qPCR of *CEBPA* expression in dTAG^V^-1 treated cells 3 days post-treatment. Fold change represents ΔΔCt values compared to the sgNT condition. n = 3 independent replicates, mean and SEM are shown. Two-sided Student t test was used for comparisons. ****p < 0.0001. **(G)** MUTZ-3 dCas9-VPR cells were infected with sgRNA-expressing lentiviruses targeting either the CEBPA cisRE or a non-targeting (NT) sequence. (Top) Histogram shows CD34 expression at day 9. (Bottom) Percentage of CD34+ cells at day 9. n = 3 independent replicates, mean and SEM are shown. **(H)** RT-qPCR of *CEBPA* expression in all conditions 3 days post-transduction. Fold change represents ΔΔCt values compared to the sgNT condition. n = 3 independent replicates, mean and SEM are shown. Two-sided Student t test was used for comparisons. ****p < 0.0001.

We next sought to characterize the dynamics of chromatin accessibility at this *CEBPA* cisRE during myeloid differentiation in primary leukemia cells (**Fig. S5D**). We divided the remission samples from the AAML1031 cohort into clusters (**Fig. S5E**) to compare chromatin accessibility within identified cisREs around *CEBPA* and approximated both *CEBPA* and *MECOM* activity along the pseudotime trajectory using gene expression and transcription factor motif enrichments from the single cell genomic data (**Fig. S5F-H**). As expected, we observed a gradual increase in both the expression and motif enrichment of CEBPA and a gradual decrease in both expression and motif enrichment of MECOM over the course of myeloid differentiation. We then investigated the accessibility of the *CEBPA* +42 kb cisRE across all defined differentiation clusters (**Fig. S5I**) and found that accessibility was generally restricted to early and later myeloid lineages (**Fig. S5J**). Together these data suggest that the cisREs regulated by MECOM are both specific and required for myeloid lineage specification. Overall, our functional screens and validation suggest a previously unappreciated and surprisingly simple regulatory logic underlying MECOM’s role in promoting stem cell-like states in AML through repression of a single critical *cis*-regulatory element.

### Repression of the *CEBPA* +42kb cisRE is sufficient to prevent myeloid differentiation induced by MECOM perturbation in primary leukemia cells

After identifying and validating the necessity for a single *CEBPA* cisRE located 42 kb downstream of the gene in aggressive stem cell-like AML cell lines, we sought to examine the necessity of this regulatory element in primary AMLs. Given the inability to create stable degron models in these sensitive, heterogeneous primary samples^46,47^, we engineered a novel CRISPR-Cas9 nuclease (Cas9n)-based strategy to disrupt *CEBPA* cisRE function. We hypothesized that a dual sgRNA approach with Cas9n could be leveraged to induce a DNA microdeletion proximal to the summit of the *CEBPA* cisRE ATAC-seq peak and corresponding MECOM ChIP-seq peak (**Fig. S6A**) to permanently inactivate the element. These sgRNAs could be co-delivered with an sgRNA targeting the coding sequence of *MECOM* to induce simultaneous inactivation of the *MECOM* locus and *CEBPA* cisRE. We initially tested this approach in MUTZ-3 MECOM-dTAG cells by electroporating them with Cas9 protein and chemically synthesized sgRNAs targeting the +42 kb *CEBPA* cisRE and *MECOM* CDS or *AAVS1* safe harbor locus. This strategy efficiently conferred a 37 bp microdeletion in the *CEBPA* cisRE in addition to creating small indels at the *MECOM* and *AAVS1* loci (**Fig. S6B**). Targeting the *MECOM* coding sequence caused near complete loss of stem cell-like CD34^+^ MUTZ-3 cells which was completely rescued by *CEBPA* cisRE inactivation (**Fig. S6C-D**). Validation of this “degron-free” Cas9n-mediated engineering strategy instilled confidence that this approach could be employed to study the relationship between MECOM and the *CEBPA* cisRE in the setting of short-term stromal cell co-cultures of these primary AML cells.

We successfully edited *MECOM* and the *CEBPA* cisRE in three primary AMLs expressing high levels of *MECOM* **(Fig. 6A-B, Table S9)**. A reduction in *MECOM* expression was confirmed in the *MECOM*-edited samples, consistent with nonsense-mediated mRNA decay^48^ (**Fig. 6C**). Moreover, across all samples *MECOM*-editing induced a significant increase in *CEBPA* expression which was almost completely prevented by the inactivation of the *CEBPA* cisRE (**Fig. 6D**). Remarkably, *MECOM* perturbations induced significant loss of stem cell-like leukemia cells as demonstrated by the loss of surface markers CD34 and/or CD117 across all patient samples (**Fig. 6E-K, Fig. S6E-F**), while inactivation of the *CEBPA* cisRE could significantly rescue this differentiation phenotype and maintain cells in more stem cell-like states. Furthermore, for one sample that grew in culture, *CEBPA* cisRE inactivation prevented the observed transient increase in cell growth following *MECOM* perturbation (**Fig. S6G**). In sum, our cisRE inactivation strategy has demonstrated the functional conservation of a key MECOM-regulated cisRE linked to *CEBPA* and demonstrated its necessity for the differentiation of primary AMLs.

**Figure 6:**
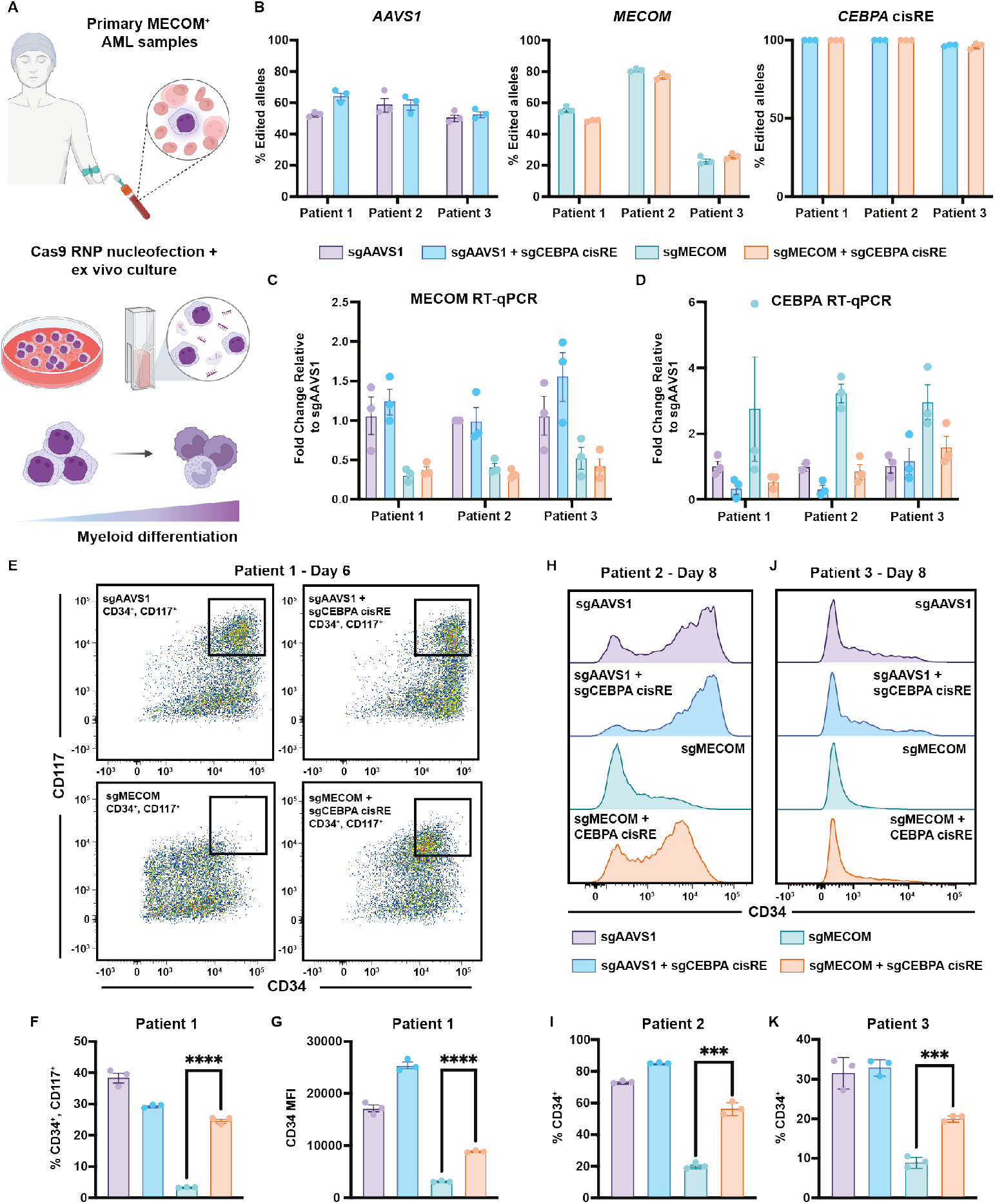
CEBPA cisRE is necessary for differentiation of MECOM-driven AML cells. **(A)** Primary MECOM+ AML cells were harvested from patients at diagnosis and cryopreserved **(Table S9)**. Cells were thawed for short-term *ex vivo* culture and electroporated with CRISPR-Cas9 RNPs to induce genetic perturbations at the *MECOM* vs. *AAVS1* locus +/−*CEBPA* +42kb cisRE. **(B)** Efficiency of gene editing in 3 biologically distinct primary AMLs at the *AAVS1, MECOM*, and *CEBPA* (cisRE) loci. Editing estimated using Sanger sequencing of amplicons followed by sequence trace decomposition analysis with ICE tool^57^. For *CEBPA* cisRE, only deletions resulting from dual guide cleavage were counted. n = 3 technical replicates, mean and SEM are shown. **(C-D)** RT-qPCR of *CEBPA* and *MECOM* expression in all conditions 3 days post-electroporation. Fold change represents ΔΔCt values compared to the sgNT condition. n = 3 technical replicates, mean and SEM are shown. **(E-G)** Immunophenotypic analysis of primary leukemia sample (patient 1, **Table S9**) 6 days post-electroporation. **(E)** Bivariate plot showing CD34 and CD117 expression assessed by flow cytometry. Black box denotes CD34+/CD117+ subset. **(F)** Percentage of CD34+/CD117+ cells. **(G)** CD34 expression measured by mean fluorescence intensity (MFI). n = 3 independent replicates, mean and SEM are shown. Twosided Student t test was used for comparisons. ****p < 0.0001. **(H-I)** Immunophenotypic analysis of primary leukemia sample (patient 2, **Table S9**) 8 days post-electroporation. **(H)** Histogram showing CD34 expression assessed by flow cytometry. **(I)** Percentage of CD34+ cells. n = 3 independent replicates, mean and SEM are shown. Two-sided Student t test was used for comparisons. ***p < 0.001. **(J-K)** Immunophenotypic analysis of primary leukemia (patient 3, **Table S9**) 8 days post-electroporation. **(J)** Histogram showing CD34 expression assessed by flow cytometry. **(K)** Percentage of CD34+ cells. n = 3 independent replicates, mean and SEM are shown. Two-sided Student t test was used for comparisons. ***p < 0.001.

### Transient activation of *CEBPA* +42kb cisRE is sufficient to promote differentiation of primary AML cells and reduce leukemic burden *in vivo*

Given the conserved role of the *CEBPA* cisRE in blocking differentiation of high-risk AMLs, we assessed whether activation of this regulatory node alone is sufficient to induce differentiation. To test this, we co-delivered *in vitro* transcribed CRISPRa (dCas9-VPR) mRNA and two chemically synthesized sgRNAs targeting the *CEBPA* cisRE or a non-targeting sgRNA into primary AML cells. Treated cells were maintained in *ex vivo* culture and monitored for signs of immunophenotypic differentiation. RT-qPCR analysis revealed that *CEBPA* cisRE targeting by CRISPRa conferred a 2-fold increase in *CEBPA* expression (**Fig. 7A**). Despite this modest activation, we observed striking differentiation phenotypes with loss of stem cell surface markers CD34 and CD117 (**Fig. 7B-D**) and an increase in expression of the mature myeloid cell marker CD11b (**Fig. 7E-F**). Notably, these differentiation phenotypes were observed across three different patient samples (**Fig. S7A-D, Table S9**). In one sample that successfully grew in culture, we also observed a marked increase in growth after CRISPRa treatment, another independent indicator of differentiation phenotypes being induced (**Fig. 7G**). To orthogonally evaluate the robustness of cell-state changes induced by *CEBPA* cisRE activation, we performed RT-qPCR analysis after 18 days of culture post-editing to assess expression of a panel of *bona fide* stem cell genes. This analysis revealed that *CEBPA* cisRE activation reduced expression of established HSC genes^38,49^ in addition to clinically relevant LSC17 genes^50^ (**Fig. 7H**). Of note, *CEBPA* cisRE activation results in reduced expression of MECOM itself showing how reactivation of myeloid differentiation is sufficient to overcome the promotion of stem cell gene expression programs.

**Figure 7:**
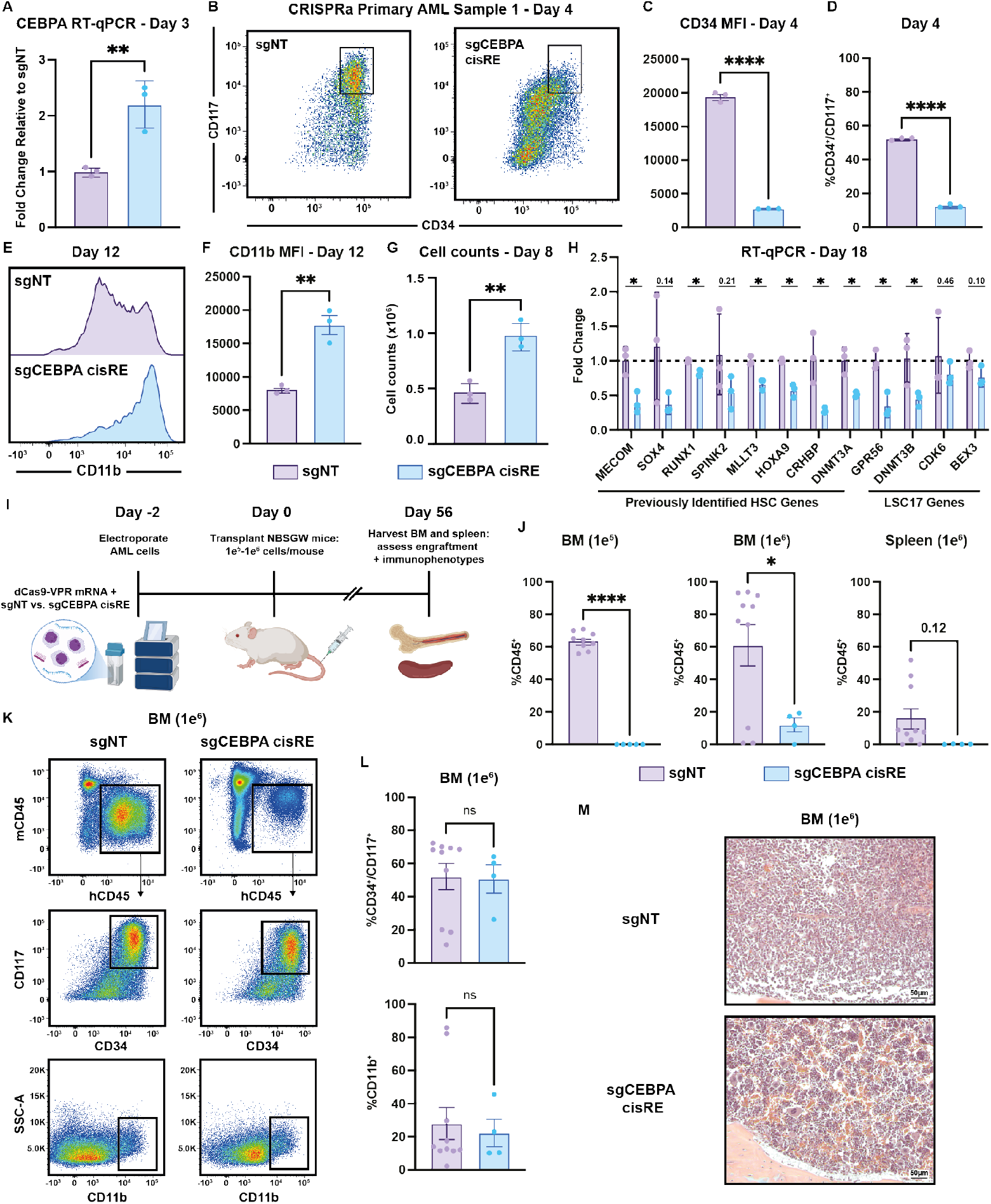
Transient activation of CEBPA cisRE is sufficient to differentiate high-risk, stem cell-like AML cells. **(A)** RT-qPCR of *CEBPA* expression 3 days post-electroporation. Fold change represents ΔΔCt values compared to the sgNT condition. n = 3 independent replicates, mean and SEM are shown. Two-sided Student t test was used for comparison. **p < 0.01. **(B-D)** Immunophenotypic analysis of primary leukemia sample (patient 1, **Table S9**) 4 days post-electroporation. **(B)** Bivariate plot showing CD34 and CD117 expression assessed by flow cytometry. Black box denotes CD34+/CD117+ subset. **(C)** CD34 expression measured by MFI. **(D)** Percentage of CD34+/CD117+ cells. n = 3 independent replicates, mean and SEM are shown. Two-sided Student t test was used for comparisons. ****p < 0.0001. **(E-F)** Immunophenotypic analysis of primary leukemia sample (patient 1, **Table S9**) 12 days post-electroporation **(E)** Histogram showing CD11b expression assessed by flow cytometry. **(F)** Percentage of CD11b+ cells. n = 3 independent replicates, mean and SEM are shown. Two-sided Student t test was used for comparison. **p < 0.01. **(G)** Viable cell counts by trypan blue exclusion in primary leukemia sample (patient 1, **Table S9**) 8 days post-electroporation. n = 3 independent replicates, mean and SEM are shown. Two-sided Student t test was used for comparison. **p < 0.01. **(H)** RT-qPCR data of a panel of established HSC genes and LSC17 genes 18 days post-electroporation demonstrating the robust differentiation of primary leukemia sample (patient 1, **Table S9**) following transient activation of *CEBPA* cisRE. n = 3 independent replicates, mean and SEM are shown. Two-sided Student t test was used for comparison. *p < 0.05. **(I)** Schematic of experiment to assess the *in vivo* impact of *CEBPA* cisRE activation of xenotransplanted primary leukemia sample (patient 1, **Table S9**). Cells were electroporated with mRNA encoding dCas9-VPR and two chemically synthesized sgRNAs targeting the *CEBPA* cisRE or a non-targeting sgRNA (sgNT). Cells recovered in *ex vivo* culture for 2 days post-electroporation and then injected via tail vein. All animals were sacrificed 56 days after transplant for analysis of leukemia burden in spleens and bone marrow. **(J)** Quantification of human cell chimerism (hCD45+) in the bone marrow of mice transplanted with 1e^5^-1e^6^ cells and spleens of mice transplanted with 1e^6^ cells. n = 4-10 xenotransplant recipients as shown, mean and SEM are shown. Two-sided Student t test was used for comparison. ****p < 0.0001, *p < 0.05. **(K-L)** Immunophenotypic analysis of the bone marrow of mice transplanted with 1e^6^ cells. Cells were labeled with cocktail of antibodies including mouse CD45, and human CD45, CD34, CD117, and CD11b. **(K)** Bivariate plots depicting gating strategy for quantification of engrafted leukemia stem/progenitor cells (CD34+/CD117+) and mature cells (CD11b+). Black boxes denote human cell subset (top), CD34+/CD117+ subset (middle), and CD11b+ subset (bottom). **(L)** Percentage of CD34+/CD117+ cells (top) and CD11b+ cells (bottom). n = 4-10 xenotransplant recipients as shown, mean and SEM are shown. Two-sided Student t test was used for comparison. ns, not significant. **(M)** Hematoxylin and eosin (H&E) staining of bone marrow of mice transplanted with 1e^6^ cells.

Finally, we performed xenotransplantation of electroporated primary AML cells into non-irradiated immuno-deficient NOD.Cg-*Kit*^W-41J^*Tyr*^+^*Prkdc*^scid^*Il2rg*^tm1Wjl^ (NBSGW) mice to assess how *CEBPA* cisRE activation impacts leukemia burden and engraftment of modified cells (**Fig. 7I**). Across two cell doses, at 8 weeks post-transplant, *CEBPA* cisRE-activated AML cells either did not engraft in any mice (1e^5^ cell dose: 0/5 mice) compared to 100% engraftment of controls (9/9 mice) or engrafted with significantly lower human chimerism in the bone marrow and spleens (1e^6^ cell dose) compared to non-targeting controls (12.1% vs. 85.5%, bone marrow hCD45^+^, p < 0.05) (**Fig. 7J, Fig. S7E**). Furthermore, the estimated leukemia initiating cell frequency, calculated based on frequencies of human cell engraftment in the bone marrow^51^ of *CEBPA* cisRE-activated AML cells, was significantly lower than non-targeting controls (1/910,241 (0.00011%) vs. 1/251,838 (0.00040%), p = 0.037) (**Fig. S7F**). We also observed an average 1.73-fold increase in spleen weights of mice transplanted with control cells compared to mice transplanted with *CEBPA* cisRE-activated cells (**Fig. S7G-H**). Notably, the lower number of cells in the *CEBPA* cisRE-activated group that did engraft in mice conferred a mostly stem cell-like immunophenotype (CD34^+^, CD117^+^, CD11b^−^) similar to those seen in controls, suggesting that these cells might escape CRISPRa activity and retain their phenotype (**Fig. 7K-L**). Analysis of fixed hematoxylin and eosin-stained bone marrow sections confirmed substantial human AML xenografts in mice transplanted with control cells (**Fig. 7M**). In contrast, mice transplanted with *CEBPA* cisRE-activated cells showed a marked reduction in leukemia burden and a high frequency of multinucleated giant cells (**Fig. 7M**). In summary, these results underscore the utility of reactivating myeloid differentiation programs in high-risk leukemia to significantly disrupt the fitness of stem cell-like leukemia cells *in vivo*.

## Discussion

A direct understanding of how stem cell gene regulatory programs are co-opted to drive aggressive AMLs is essential for developing targeted therapeutic strategies. Here, we have focused on deciphering the precise mechanisms enabling such stem cell gene regulatory programs upon elevated MECOM expression, as is seen in many high-risk AMLs. Motivated by clinical observations, including the recent occurrence of MECOM insertions in driving aggressive leukemias in gene therapy trials^52^, we utilized targeted MECOM degradation in conjunction with functional genomic readouts to define highly conserved gene and corresponding *cis*-regulatory element networks that are directly repressed by MECOM. The identification of these networks highlights the primary role of MECOM in AML as a strong transcriptional repressor that suppresses differentiation programs, a role that was previously underappreciated amongst a multitude of other proposed functions^15–19^.

We sought to understand this role through functional genomic perturbations and high throughput screens to identify key nodes within this direct MECOM network that play a pivotal role in enabling its ability to maintain leukemia cells in stem cell-like states. By focusing on MECOM-regulated chromatin sites, rather than genes themselves, we hypothesized that we could more accurately recapitulate or reverse MECOM’s endogenous gene regulatory activity. Remarkably and unexpectedly, within this vast network, we observed that modulation of a single *cis*-regulatory element of *CEBPA*, is both necessary and sufficient to sustain MECOM’s role in repressing myeloid differentiation in aggressive AMLs. This is surprising given the hundreds of chromatin sites under strong regulation by MECOM and highlights the importance of utilizing functional screens to interrogate gene regulatory networks to identify therapeutic vulnerabilities.

The key functional relationship we have elucidated between MECOM and CEBPA is intriguing given the disparate clinical outcomes of unfavorable MECOM^+^ AMLs and more favorable CEBPA-mutant AMLs^53,54^. However, when stratifying AMLs by mutations in one vs. both copies of *CEBPA* across many cohorts, single mutant CEBPA AML confers significantly worse patient outcomes^55,56^. This is consistent with our observation of MECOM’s role in stem cell-like leukemias, where MECOM regulates CEBPA not through complete repression, but rather by allowing a residual basal level of CEBPA expression and function in stem cell-like states in AMLs that express *MECOM*. We also cannot rule out the possibility that differences in the cell of origin of oncogenic transformation contribute to the potentially distinct cell states observed in *MECOM* expressing versus *CEBPA*-mutated leukemia cells, and their differing clinical outcomes.

This work establishes proof-of-concept for an approach that enables differentiation of high-risk AMLs driven by MECOM by simply activating a single differentiation factor, CEBPA. Current strategies that seek to eradicate stem cell-like populations in AML, and which have proven of limited effectiveness, might not be ideal approaches for these incurable leukemias. Rather, the paradigm established in acute promyelocytic leukemia using differentiation therapy, could also be successfully applied in high-risk AMLs by reactivating CEBPA expression or function.

## Acknowledgements

We are grateful to members of the Sankaran laboratory for valuable discussions and feedback. We also thank the Boston Children’s Hospital Viral Core for assistance with rAAV production. This work was supported by the New York Stem Cell Foundation (V.G.S.), the Howard Hughes Medical Institute (V.G.S.), the Alex’s Lemonade Stand Foundation (V.G.S.), and National Institutes of Health (NIH) grants R01CA265726, R01CA292941, R33CA278393, R01DK103794, and R01HL146500 (V.G.S.), as well as R35 CA283977 (K.S.). T.J.F. received support from the NIH grant F31CA287658 and the National Science Foundation Graduate Research Fellowship. R.A.V. received support from NIH grant K08 CA286756, the Edward P. Evans Foundation, the Alex’s Lemonade Stand Foundation, the Cancer Prevention and Research Institute of Texas, and is a Horchow Family Scholar. V.G.S. is an Investigator of the Howard Hughes Medical Institute.

## Author contributions

T.J.F, M.C.G, R.A.V., and V.G.S. conceptualized the study. T.J.F, S.L., M.C.G, R.A.V., and V.G.S. devised the methodology. T.J.F., M.A., S.L., M.C.G., R.P., L.W., S.S., F.E.R., K.D.D, J.A.P., C.M., M.N.B., and R.A.V. performed studies. T.J.F., M.A., S.L., C.M., K.A., and V.G.S. provided data visualization media. A.A., M.D.M, J.A.P., Y.P., and K.S., provided primary AML samples. K.A., K.S., K.R.M., S.P.G., V.H., M.D.M., contributed ideas and insights. V.G.S. acquired funding for this work and provided overall project oversight. T.J.F. and V.G.S. wrote the original manuscript, as well as edited the manuscript with input from all authors.

## Data and code availability

Raw and processed data have been deposited at GEO and will be available at the time of publication.

## Materials and Methods

### Data reporting

No statistical methods were used to predetermine sample sizes but our sample sizes are similar to those reported in previous publications^49,58^. Data collection and analysis were not performed blind to the conditions of the experiments. No animals or data points were excluded from analysis.

### Cell line and primary AML cell culture

MUTZ-3 cells (DSMZ), HNT-34 cells (Creative Bioarray), and OCI-AML4 cells (DSMZ) were cultured at 37 °C in α-MEM (Life Technologies) supplemented with 20% FBS, 20% conditioned medium from 5637 cells^59^ (ATCC) and 1% penicillin/streptomycin. Confluency for all cells was maintained between 7 × 10^5^ and 1.5 × 10^6^ ml^−1^.

UCSD-AML1 cells (a gift from Dr. Kimberly Stegmaier’s lab) were cultured at 37 °C in RPMI 1640 (Life Technologies) supplemented with 20% FBS, 1% penicillin/streptomycin and 10ng/mL GM-CSF (Peprotech). Confluency was maintained between 5 × 10^5^ and 1.5 × 10^6^ ml^−1^.293T cells were cultured at 37 °C in DMEM (Life Technologies) supplemented with 10% FBS and 1% penicillin/streptomycin.MS-5 cells, prior to co-culture, were cultured at 37 °C in α-MEM (Life Technologies) supplemented with 10% FBS, 2 mM L-glutamine and 2 mM sodium pyruvate. Confluency was maintained at <95% and split in culture 1:3 every 3 days. Cells were maintained at low passage number (<12).

Primary AML cells were collected with informed consent according to procedures approved by either the University Health Network (UHN) or Boston Children’s Hospital and Dana Farber Cancer Institute’s Research Ethics Boards. Two days prior to thawing primary AML cells, MS-5 cells were plated at ~50% confluency in 12-well or 6-well plates. Primary AML cells were then thawed and immediately placed in co-culture with MS-5 cells. Cells were co-cultured at 37 °C in IMDM (Life Technologies) supplemented with 10% FBS, 2% L-glutamine, CC100 cytokine cocktail (Stem Cell Technologies), and 100 ng/ml TPO at concentrations between 1 and 2.5 × 10^6^ ml^−1^.

### Mouse model

NOD.Cg-*Kit*^W-41J^*Tyr*^+^*Prkdc*^scid^*Il2rg*^tm1Wjl^ (NBSGW) mice were obtained from the Jackson Laboratory (stock 026622)^60^. Littermates of the same sex were randomly assigned to experimental groups. NBSGW were interbred to maintain a colony of animals homozygous or hemizygous for all mutations of interest. The Institutional Animal Care and Use Committee at Boston Children’s Hospital approved the study protocol and provided guidance and ethical oversight.

### Lentiviral production

For lentiviral production, 293T cells were expanded to reach 80% confluency per plate on the day of transfection. 1–20 10 cm^2^ plates were prepared per lentiviral construct. For each plate, 4 µg of psPAX2 packaging plasmid, 2 µg of pMD2.G envelope plasmid, and 8 µg of sgRNA vector construct was mixed in Opti-MEM media (Gibco, 31985-062). This mix was then diluted in Lipofectamine 3000 and combined with P3000 reagent per the manufacturer’s protocol and added dropwise to cells. 12-16 hours later, 293T medium was removed and changed to DMEM with 20% FBS and 1% penicillin/streptomycin. 24 hours later the media was harvested and filtered through a Stericup 0.45 mm PVDF membrane (Millipore, SCHVU01RE), and transferred to ultra-clear centrifuge tubes (Beckman Coulter, 344058). Virus was subsequently concentrated using a Beckman Coulter SW32Ti

Ultracentrifuge with the following parameters: Speed: 24,000 rpm, time: 1 hour and 30 minutes, Temperature: 4C, maximum acceleration and deceleration 9. The supernatant was removed, and the virus pellet was resuspended with the appropriate media. Concentrated virus was stored at −80ºC until further usage.

### Lentiviral transduction

Cells were transduced at a density of 500,000-1 million cells per mL. Concentrated virus was added to cells along with 8 µg/mL polybrene (Sigma Aldrich, TR-1003-G). Cells were then spinfected at 2,000 rpm for 90 mins at 37ºC. 12-16 hours after spinfection, the media was replaced by the appropriate complete media.

### Transplantation assays

Primary AML cells were thawed and plated onto an MS-5 co-culture with primary cell medium (see methods) to recover for 24 hours. Cells were then electroporated and placed back into co-culture. 48 hours later, non-irradiated NBSGW mice (between 8-10 weeks of age) were tail vein injected with modified primary AML cells (1 × 105 - 1 × 106 cells). Peripheral human chimerism was assessed at 4 and 7 weeks and animals were sacrificed at 8 weeks for bone marrow (BM) and spleen. Human chimerism and corresponding immunophenotypes were assessed by flow cytometry. The relative percentages of human chimerism and cell counts were used in conjunction to quantify the size (cellular quantity) of human leukemia cell xenografts in the BMs and spleens. Functional leukemia initiating cell (LIC) frequencies were calculated based on human cell engraftment frequencies in the bone marrow of transplanted mice using the ELDA software51. Engraftment of leukemia cells was considered to have occurred if human cell chimerism in the bone marrow was >5%. A piece of spleen and a whole femur from each mouse was fixed in Bouin’s fixative solution for over 24 hours, followed by two consecutive washes in 70% ethanol. Subsequently, bones were decalcified with formic acid, samples were embedded in paraffin, 4 µm sectioned and H&E stained at the Rodent Histopathology Core at Harvard Medical School. Stained slides were analyzed on Zeiss Axio Imager Z2 Microscope at the Cellular Imaging Core Facility Boston Children’s Hospital.

### Flow cytometry and cell sorting

MUTZ-3 and UCSD-AML1 cells were stained with anti-CD34-APC (BioLegend, 343607). MUTZ-3 cells were also stained with anti-CD14-PE-Cy7 (BioLegend, 367112). *Ex-vivo* cultured Primary AML samples for were stained with anti-CD34-Alexa Fluor® 488 (BioLegend, 343518), anti-CD117-PE (BioLegend, 313204), and/or anti-CD11b-PE-Cy7 (BioLegend, 101216). Xenotransplant samples were stained with anti-human CD34-BV421 (BioLegend, 343610), anti-human CD117-PE (BioLegend, 313204), antihuman CD11b-PE-Cy7 (BioLegend, 101216), anti-human CD45-APC (BioLegend, 304037), and anti-mouse CD45-FITC (BioLegend, 103108). Two microliters of each antibody were used per 1 × 10^5^ cells in 100 µl in all experiments.

Flow cytometric analyses were conducted on a BD LSRII, LSR Fortessa or Accuri C6 instruments and all data were analyzed using FlowJo software (v.10.10). FACS was performed on a BD Aria in a sterile biosafety cabinet where samples were collected in PBS containing 2% FBS and subsequently replated in the appropriate human cell culture medium. Alternatively, for molecular analyses of sorted populations (RT-qPCR or gDNA PCR for CRISPR screens) cells were sorted into Eppendorf or conical tubes containing Buffer RLT Plus (QIAGEN) with 1% BME and immediately frozen at −80 °C for downstream analyses.

### Cell cycle analysis

MUTZ-3-dTAG and UCSD-AML1-dTAG cells were treated with 500nM dTAG^V^-1 or DMSO for 48 hours, incubated with 5-ethynyl-2′-deoxyuridine (EdU) (Thermo Fisher Scientific, C10636), then stained with a fluorescent CD34 antibody. Cells were then fixed, permeabilized per the manufacturer’s recommendations and incubated with propidium iodide (PI) to stain total DNA content. Flow cytometry analysis ran at low speeds (<400 events/second) was then performed to assess cell cycle states.

### Western blot analysis

Total protein lysate of cells was extracted by RIPA buffer in presence of protease inhibitor cocktail on ice for 30 minutes. The total lysate was then linearized by 1X SDS loading buffer and heated at 55 °C for 10 minutes. The lysate was loaded onto 4-20% Mini-PROTEAN TGX Precast Protein Gels (BioRad, 456104) before being transferred to a PVDF membrane using BioRad Trans-Blot Turbo Transfer System. The membrane was blocked in LI-COR Intercept Blocking Buffer and incubated with primary antibodies at 1:1000 dilution in LI-COR Intercept Antibody Diluent at 4 °C overnight, then washed and incubated in 1:1000 anti-mouse-HRP secondary antibody for 90 minutes at room temperature. Membrane was developed using BioRad Clarity Western ECL Substrate and Reagent (BioRad, 1705061) and imaged with BioRad system.

### Bulk RNA-seq

Total RNA was extracted using the RNeasy Micro kit (QIAGEN, 74004) from 10,000-50,000 cells sorted or harvested in 25 µl Buffer RLT Plus with 1% BME. Then we proceeded with the SmartSeq2 protocol from the reverse transcription step using 10 ng of RNA^61^. The whole transcriptome amplification step was set at ten cycles. Bulk RNA libraries were pooled at appropriate molar ratios to obtain at least 20 million reads per library. Libraries were subject to paired-end sequencing using NextSeq550 High Output kits with 150 cycles.

### ATAC-seq

Accessible chromatin was assessed using Assay for Transposase-Accessible Chromatin using sequencing (ATAC-seq) as previously described with the Omni-ATAC protocol^62^ with minor adaptations detailed here. *MECOM-FKBP12*^*F36V*^ modified MUTZ-3, UCSD-AML1, HNT-34 and OCI-AML4 cells were treated with 500nM dTAG^V^-1 or DMSO for 6 hours and then 50,000 live cells were sorted into PBS with 2% BSA. Cells were washed twice in 150µL and 50µL 1xPBS, resuspended in 50µL ATAC-seq lysis buffer, incubated for 10 min on ice and centrifuged at 400g for 10 min at 4 °C. The pellet was incubated in the transposase reaction mix (25µL 2×TD buffer (Illumina), 2.5µL transposase (Illumina Cat# FC-121-1030) and 22.5µL nuclease-free water) for 30min at 37 °C with gentle agitation. After DNA purification with the Zymo DNA Clean and Concentrator Kit (Zymo, Cat# D4033) libraries were amplified with NEBNext High-Fidelity PCR Master Mix (NEB, Cat# M0541S) using custom Nextera primers. Libraries for sequencing were size selected with Agencourt AMPure XP beads (Beckman Coulter, Cat# A63880). DNA concentration was measured with an Invitrogen Qubit fluorometer (Life Technologies) and Agilent Fragment Analyzer. The libraries were sequenced using the Illumina NextSeq 500 platform and the 75-bp paired-end configuration to obtain at least 30 million reads per sample.

### PRO-seq

Aliquots of frozen (−80C) permeabilized MUTZ-3 cells were thawed on ice and pipetted gently to fully resuspend. Aliquots were removed and permeabilized cells were counted using a Luna II, Logos Biosystems instrument. For each sample, 1 million permeabilized cells were used for nuclear run-on, with 50,000 permeabilized Drosophila S2 cells added to each sample for normalization. Nuclear run on assays and library preparation were performed essentially as described in Reimer et al. 2021^63^ with modifications noted: 2X nuclear run-on buffer consisted of (10 mM Tris (pH 8), 10 mM MgCl2, 1 mM DTT, 300mM KCl, 20uM/ea biotin-11-NTPs (Perkin Elmer), 0.8U/uL SuperaseIN (Thermo), 1% sarkosyl). Run-on reactions were performed at 37C. Adenylated 3’ adapter was prepared using the 5’ DNA adenylation kit (NEB) and ligated using T4 RNA ligase 2, truncated KQ (NEB, per manufacturer’s instructions with 15% PEG-8000 final) and incubated at 16C overnight. 180uL of betaine blocking buffer (1.42g of betaine brought to 10mL with binding buffer supplemented to 0.6 uM blocking oligo (TCCGACGATCCCACGTTCCCGTGG/3InvdT/)) was mixed with ligations and incubated 5 min at 65C and 2 min on ice prior to addition of streptavidin beads. After T4 polynucleotide kinase (NEB) treatment, beads were washed once each with high salt, low salt, and blocking oligo wash (0.25X T4 RNA ligase buffer (NEB), 0.3uM blocking oligo) solutions and resuspended in 5’ adapter mix (10 pmol 5’ adapter, 30 pmol blocking oligo, water). 5’ adapter ligation was per Reimer et al. 2021 but with 15% PEG-8000 final. Eluted cDNA was amplified 5-cycles (NEBNext Ultra II Q5 master mix (NEB) with Illumina TruSeq PCR primers RP-1 and RPI-X) following the manufacturer’s suggested cycling protocol for library construction. A portion of preCR was serially diluted and for test amplification to determine optimal amplification of final libraries. Pooled libraries were sequenced using the Illumina NovaSeq platform.

### ChIP-seq

Chromatin immunoprecipitation followed by sequencing (ChIP-seq) was performed on chromatin from 1×10^6^ CD34^+^ MUTZ-3-dTAG cells after treatment with 500nM dTAG^V^-1 or DMSO for 6 hours. Cells were cross-linked with 1% methanol-free formaldehyde (Pierce Life Technologies, 28906), quenched with 0.125 M glycine and frozen at −80 °C and stored until further processing. ChIP reaction was performed with iDeal ChIP-seq kit for TFs (Diagenode, C01010055) with modifications of the manual detailed below. Lysed samples were sonicated using the E220 sonicator (Covaris, 500239) in microTUBE AFA Fiber Pre-Slit Snap-Cap tubes (Covaris, 520045) with settings for 200-bp DNA shearing. Sheared chromatin was immunoprecipitated with 2.5 µg HA antibody (CST, HA-Tag (C29F4) Rabbit mAb #3724), 2.5 µg H3K27ac antibody (Diagenode, C15410196) or 2.5 µg IgG antibody (Diagenode, C15410206, RRID AB_2722554). Eluted and decross-linked DNA was purified with MicroChIP DiaPure columns (Diagenode, C03040001) and eluted in 30 µl of nuclease-free water. ChIP and input libraries for sequencing were prepared with ThruPLEX DNA-Seq kit (Takara, R400674) and DNA Single Index kit, 12S Set A (Takara, R400695). Size selection steps were performed with Magbio Genomics HighPrep PCR beads (Fisher Scientific, 50-165-6582). The libraries were sequenced using the Illumina NextSeq 500 platform and the 150-bp paired-end configuration to obtain at least 20 million reads per sample.

### Annexin V apoptosis staining

HNT-34-dTAG cells were treated with 500nM dTAG^V^-1 or DMSO in culture for three days and then collected. Cells were washed twice with cold PBS with 2% FBS and then resuspended in Annexin V Binding Buffer (Cat. No. 422201) at a concentration of 1×10^6^ cells/ml. 100 µl of cell suspension was transferred to an Eppendorf tube and mixed with 5 µl of APC Annexin V. Propidium iodide was added then cells were gently vortexed and incubated for 15 min at room temperature (25°C), in the dark. 400 µl of Annexin V Binding Buffer was then added to each sample and cells were analyzed by flow cytometry.

### CRISPR engineering of cell lines and primary AML cells

Cell lines were electroporated using the Lonza 4D Nucleofector with 20 µl Nucleocuvette strips as described^49,64^. Cas9-sgRNA ribonucleoprotein complexes (RNPs) were made by combining 50 pmol of Cas9 protein (IDT) and 100 pmol of chemically synthesized sgRNA (Synthego) targeting the C-terminus of *MECOM* and incubating at 21 °C for 15 min. Between 2 × 10^5^ and 5 × 10^5^ cells were resuspended in 20 µl P3 solution, mixed with RNP and underwent nucleofection with program EO-100. Cells were returned to appropriate cell culture medium and supplemented with 50µl of crude rAAV HDR donor (see methods) and 1µM of alt-R HDR enhancer (IDT, 10007910). 24 hours later, cells were washed once with 1x PBS and replated in fresh cell culture medium. 72 hr after electroporation cells were analyzed via flow cytometry to assess GFP expression. Most AML models we chose, including MUTZ-3, UCSD-AML1, and HNT-34, exhibit a translocation or inversion on chromosome 3^22,23^ which hyperactivates expression of a single copy of the *MECOM* locus. This allowed for sorting of polygenic populations of GFP^+^ cells in which the transactivated *MECOM* allele was correctly tagged with the *FKBP12*^*F36V*^ degron cassette. On-target editing and successful knock-in was further confirmed by genomic DNA PCR. Genomic DNA was extracted using the DNeasy kit (QIAGEN) according to the manufacturer’s instructions. PCR was performed using Platinum II Hotstart Mastermix (Thermo Fisher Scientific) and primers flanking the repair site. For OCI-AML4 cells, which do not have an inversion or translocation on chromosome 3 but still exhibit relatively high MECOM expression, following CRISPR editing single cells were plated by limiting dilution in 96-well plates and clonally expanded. After 2 weeks of expansion, genomic DNA was extracted from clones and screened via PCR for biallelic tagging. A homozygous clone was identified and further expanded in culture for experimental use.

Primary AML cells were electroporated using the Lonza 4D Nucleofector with 20 µl Nucleocuvette strips. For Cas9 nuclease experiments, 50 pmol of Cas9 protein was mixed with a total 100 pmol of sgRNAs targeting either the *MECOM* or *AAVS1* loci alone or with sgRNAs targeting the CEBPA cisRE. For CRISPRa mRNA experiments, 2.5 µg of dCas9-VPR mRNA was mixed with 100 pmol of sgRNAs targeting the CEBPA cisRE or a non-targeting control. In all experiments between 2 × 10^5^ and 5 × 10^5^ cells were resuspended in 20 µl P3 solution, mixed with the corresponding CRISPR reagents, and underwent nucleofection with program DZ-100. Electroporated cells were maintained in MS-5 co-culture with the appropriate medium. To assess editing efficiencies in Cas9 nuclease experiments, genomic DNA PCR was performed using Platinum II Hotstart Mastermix (Thermo Fisher Scientific) and edited allele frequency was detected by Sanger sequencing and analyzed by ICE^57^ (Table S1-2). The effect of CRISPR editing on gene expression was assessed by RT-qPCR three days after electroporation.

### Recombinant AAV production

The triple-transfection method was used to generate crude rAAV lysates^65^. HEK293T cells were plated in a 10 cm dish in 1X Penicillin-streptomycin-Glutamine and 10% FBS in DMEM. At 80% confluence, the medium was replaced with fresh medium. Then, cells were triple transfected using polyethylenimine Max (PEI Max, Polysciences 24765-1) with 12 µg of pAAVhelper, 7.5 µg of pRep2Cap6, and 7.5 µg of transfer plasmid (PEI:DNA = 3:1), in DMEM without phenol red (Life Technologies 31053036). 3 days after transfection, cells were scraped and collected by spinning at 1300 RPM for 5 min. Cell pellet was resuspended in 1-2 mL DPBS without calcium or magnesium and lysed by three rounds of freeze-thaw. This was accomplished by placing them alternately in a dry ice/ethanol bath until completely frozen and in a water bath of 37 °C until completely thawed. After the final thaw, cell lysate was spun at 1300 RPM for 5 min and the supernatant was filtered through a 0.22 µm syringe filter to yield a crude viral lysate. Viral lysates were stored in 25 µL aliquots at 4C for up to 4 weeks, and at −80C thereafter.

### RNA isolation, reverse transcription, and real-time PCR

RNA was harvested by the RNeasy Micro kit (QIAGEN, 74004) and quantified by nanodrop. 1 microgram of total RNA was then reverse transcribed using iScript cDNA synthesis kit following manufacturer’s instructions. The reverse transcribed cDNA was then diluted (1:20) and real time PCR was run using Biorad iQ SYBR green supermix. Data was normalized by loading control (*ACTB*) and presented as fold change compared to control samples using the delta-delta CT (ΔΔCT) method.

### Quantitative mass spectrometry-based proteomics

MUTZ-3 cells were treated with DMSO or 500 nM dTAGV-1 and lysed in a buffer containing 8 M urea and 200 mM EPPS at pH 8.5 with protease inhibitors. The lysates were generated using a probe sonicator (20 pulses of 0.5 seconds at level 3). Protein concentration was measured using a BCA assay, and 50 µg of protein was aliquoted for each condition. Proteins were reduced with TCEP for 15 minutes at room temperature (RT) and alkylated with 10 mM iodoacetamide for 30 minutes in the dark at RT. Precipitation was performed using chloroform/methanol, as described previously^66^. Samples were digested overnight with LysC and trypsin (1:100 enzyme/protein ratio) at 37°C on a ThermoMixer set to 1,200 rpm. After digestion, peptides were labeled with TMTpro 18-plex reagents (1:2 peptide/reagent mass ratio) for 1 hour with constant shaking at 1,200 rpm. Excess TMT reagent was quenched with 0.3% hydroxylamine for 15 minutes at RT. The samples were mixed in equal proportions across all TMT channels, pooled, and dried using a Speedvac.

The pooled peptides were desalted with a 100-mg Sep-Pak solid-phase extraction cartridge. After desalting, the peptides were dried, resuspended in a buffer (10 mM ammonium bicarbonate, 5% acetonitrile, pH 8.0), and fractionated into a 96-well plate using basic pH reversed-phase HPLC with an Agilent 300 Extend-C18 column. Fractionation was performed with a 50-minute linear gradient of 13–43% buffer (10 mM ammonium bicarbonate, 90% acetonitrile, pH 8.0) at a flow rate of 0.25 mL/min. The peptide mixture was combined into 24 fractions and were desalted using StageTips^66^. Forty percent of the resuspended sample (10 µL of 5% acetonitrile, 5% FA) was analyzed on an Orbitrap Eclipse using a high-resolution MS2-based method.

### Liquid chromatography and mass spectrometry data acquisition

Mass spectrometry data were collected using a Orbitrap Eclipse mass spectrometer (Thermo Fisher Scientific, San Jose, CA) coupled with Neo Vanquish liquid chromatograph. Peptides were separated on a 100 µm inner diameter microcapillary column packed with ~35cm of Accucore C18 resin (2.6 µm, 150 Å, Thermo Fisher Scientific). For each analysis, we loaded ~2 µg onto the column. Peptides were separated using a 90 min gradient of 5 to 29% acetonitrile in 0.125% formic acid with a flow rate of 400 nL/min. The scan sequence began with an Orbitrap MS^1^ spectrum with the following parameters: resolution 60K, scan range 350-1350, automatic gain control (AGC) target 100%, maximum injection time “auto,” and centroid spectrum data type. We use a cycle time of 1s for MS^2^ analysis which consisted of HCD high-energy collision dissociation with the following parameters: resolution 50K, AGC 200%, maximum injection time 86ms, isolation window 0.6 Th, normalized collision energy (NCE) 36%, and centroid spectrum data type.

Dynamic exclusion was set to automatic. The FAIMS compensation voltages (CV) were −40, −60, and −80V.

### CRISPR library and individual sgRNA cloning

The following protocol was used for creating a sgRNA lentiviral library or individual sgRNA lentiviruses. The sgRNA library for both screens was designed to target a conserved network of MECOM-regulated cisREs. We first mined the ENCODE Consortium’s recently published repository of functionally validated sgRNA sequences^67^ that overlapped our genomic regions of interest and selected 5 sgRNA sequences per region. For regions absent from the ENCODE database or corresponding to less than 5 validated sgRNAs we utilized the CRISPick tool from The Broad Institute to design additional sgRNAs. Oligonucleotide pools for CRISPR screens were ordered from IDT at a 50 pmol scale (standard desalting) and resuspended at a 10 µM concentration (**Table S8**). Single oligonucleotides for individual sgRNA lentiviruses were ordered from Azenta Life Sciences at a 25 nmol scale (standard desalting) and resuspended at a 10 µM concentration.

An initial extension reaction was performed using the oligo pool (**Table S8**) or individual oligonucleotides, NEB Q5 Hot Start High-Fidelity 2X Master Mix (M0492L) and extension primers. The following parameters were used for extension: 98ºC for 2 minutes; 10 cycles of (64ºC for 30 seconds and 72ºC for 20 seconds); 72ºC for 2 minutes; and hold at 4ºC. The product was purified using the Monarch® PCR & DNA Cleanup Kit (NEB, T1030), and eluted in 25 mL of water. All sgRNAs were cloned into a modified CROP-seq-opti vector^68^ in which the the puromycin resistance cassette was replaced with tag red fluorescent protein (tag-RFP) to facilitate lentiviral titration and precise FACS-enrichment of infected cells. BsiWI (NEB) and MluI (NEB) were used to excise the puromycin resistance marker and a gBlock with the tag-RFP sequence was cloned into the digested vector using the same restriction site overhangs. This modified vector was digested at 37ºC for 1 hour and purified with Monarch® DNA Gel Extraction Kit (NEB #T1020).

An NEBuilder HiFi DNA Assembly (NEB, #E2621) reaction was performed using 500 fmol of vector and 10,000 fmol of purified extension reaction product (with the volume required for each calculated using its fragment length and its concentration measured by Nanodrop) with 10 µl of 2x NEBuilder HiFi DNA Assembly Master Mix and nuclease-free water to a final reaction volume of 20 µl. The reaction was incubated at 37ºC for 2 hours. For library cloning, 5 µl of crude NEBuilder HiFi DNA Assembly product were transformed into Endura Electrocompetent Cells (Biosearch Technologies, 71003-038) using the Biorad Gene Pulser Xcell Total Electroporation System (1652660) with the following parameters: 1.8 kV, 25 µF and 200 U. Bacteria were recovered for 20 minutes in the kit’s recovery media. 2 µL of bacteria were used to create 4 serial dilutions to evaluate the transformation efficiency (and ensure at least 100x coverage of the library) and the remaining bacteria were inoculated in 500 mL of LB with 100 µg/ml of ampicillin and grown overnight at 30ºC. 16-18 hours later, plasmid DNA was extracted using the NucleoBond Xtra Maxi kit for endotoxin-free plasmid DNA (Macherey-Nagel, 740424.50) and eluted in 400 µL of nuclease-free water. For single sgRNA cloning, 2 uL of crude NEBuilder HiFi DNA Assembly product were transformed into chemically competent NEB® 10-beta Competent *E. coli* per the manufacturer’s recommended protocol and plated onto LB agar plates with 100 µg/ml of ampicillin and grown overnight at 37ºC. The following day, colonies were picked, expanded overnight at 37ºC, and plasmid DNA was extracted using the Monarch® Plasmid Miniprep Kit and eluted in 30 µL of nuclease-free water. Plasmid DNA was then sequenced by whole plasmid sequencing (Primordium) to identify positive clones.

### MECOM lentiviral overexpression constructs

The EVI1 isoform of the *MECOM* locus was used for all lentiviral overexpression experiments. A construct containing the EVI1 coding sequence and IRES-eGFP cassette placed downstream from a constitutive promoter (HIV/MSCV hybrid LTR)^69^ were obtained from Voit et al. 2023^14^. For dTAG^V^-1 rescue experiments, the IRES-eGFP cassette was first replaced with IRES-TagRFP using EcoRI and PacI restriction sites. The PLASS mutation was introduced into the PLDLS motif of the EVI1 coding sequence using two PstI restriction sites flanking the PLDLS motif. This small fragment was excised and replaced with an identical sequence, except for the PLDLS motif, which was altered to PLASS.

### CRISPRi screen

MUTZ-3-dTAG cells were transduced at high MOI with separate lentiviruses packaged with TRE-KRAB-dCas9-IRES-BFP^70^ and pLVX-EF1alpha-Tet3G (Takara #631359). TRE-KRAB-dCas9-IRES-BFP was a gift from Eric Lander (Addgene plasmid # 85449; http://n2t.net/addgene:85449; RRID:Addgene_85449). The next day, to bypass G418 selection and select for co-transduced cells, 1µg/ml of doxycycline was added and 48 hours later BFP^+^ cells were sorted by FACS. These MUTZ-3 cells stabling expressing inducible dCas9-KRAB were then transduced with a MECOM-regulated cisRE-targeting lentiviral sgRNA library at a low MOI (<0.3) in technical triplicate at ~2000x coverage (cells/sgRNA). 24 hours after transduction, 1µg/ml doxycycline was added to induce dCas9-KRAB expression and 48 hours after transduction 500nM of dTAG^V^-1 was added to all replicates. Cells were maintained in culture with regular media changes and supplementation of fresh doxycycline and dTAG^V^-1 every three days. A population of cells not treated with dTAG^V^-1 or not transduced with the sgRNA library were also maintained as controls to ensure dTAG^V^-1 treatment induced robust MUTZ-3 differentiation and that expression of the sgRNA library resulted in a relative enrichment of CD34^+^ stem-like cells, respectively (**Fig. S5A**). 14 days post-transduction, residual CD34^+^ stem-like cells were sorted by FACS and genomic DNA was extracted with the DNeasy kit (QIAGEN) according to the manufacturer’s instructions. PCR on genomic DNA from all replicates was performed using Titanium Taq DNA Polymerase and PCR buffer (Clontech Takara Cat# 639208) according to The Broad Institute Genetic Perturbation Platform’s protocol for PCR of sgRNAs from genomic DNA for Illumina sequencing (https://portals.broadinstitute.org/gpp/public/resources/protocols). In brief, barcoded P5 and P7 PCR primers (table) were used to amplify the sgRNA spacer sequence from the genomically-integrated lentiviral sgRNA expression cassette. An aliquot of sgRNA library plasmid DNA was also used as a template for this PCR to assess the coverage of the cloned sgRNA library and generate a baseline sgRNA distribution to assess relative enrichment or depletion in our screens. Amplicon libraries were purified with AMPure XP-PCR magnetic beads, and pooled at an equimolar concentration. Libraries were subject to paired-end sequencing using NextSeq550 High Output kits with 75 cycles to ensure at least 20 million reads per library.

### CRISPRa screen

MUTZ-3-dTAG cells were transduced at a high MOI with lentivirus packaged with pXPR_120 (dCas9-VPR-2A-BlastR)^71^. pXPR_120 was a gift from John Doench & David Root (Addgene plasmid # 96917; http://n2t.net/addgene:96917; RRID: Addgene_96917). 48 hours after transduction, cells were treated with 10 µg/ml of Blasticidin for 7 days to select for cells stably and constitutively expressing dCas9-VPR. These MUTZ-3 cells were then transduced with a MECOM-regulated cisRE-targeting lentiviral sgRNA library at a low MOI (~0.33) in technical triplicate at ~2000x coverage (cells/sgRNA). Cells were maintained in culture with regular media changes every three days. 14 days post-transduction, cells in the bottom 5% of CD34 expression measured via flow-cytometry were sorted by FACS. Genomic DNA extraction, library preparation, and sequencing of CRISPRa libraries was all performed as previously described for the CRISPRi screen.

### In vitro transcription of dCas9-VPR mRNA

An in vitro transcription template encoding dSpCas9-VPR was a gift from Rasmus Bak (Addgene plasmid # 205247; http://n2t.net/addgene:205247; RRID:Addgene_205247)^72^. The plasmid was digested with SapI (NEB), a restriction site immediately downstream from the encoded polyA tail, for 1 hr at 37ºC and the linear transcription template was purified using Monarch® DNA Gel Extraction Kit and eluted in 20 uL of nuclease free water. dCas9-VPR mRNA was transcribed from this template using the HiScribe T7 High Yield RNA Synthesis Kit (New England Biolabs) according to the manufacturer’s recommended protocol for high-yield synthesis with the following changes: UTP was fully replaced with N1-methylpseudouridine-5′-triphosphate (TriLink Biotechnologies) and co-transcriptional capping by CleanCap Reagent AG (TriLink Biotechnologies) was used at a ratio of 4:1 with GTP. mRNA products were precipitated in 2.5 M lithium chloride, washed twice with 70% ethanol, dissolved in nuclease-free water, and stored at −80 °C.

### AML cell line bulk RNA-seq analysis

FASTQ files were demultiplexed with bcl2fastq and aligned to the hg38 reference genome using bowtie2 (version 2.5.2). Sam files were sorted, indexed, and converted to bam files with Samtools (version 1.18). For data visualization in a genome browser, bam files were converted to bigwig files using the bamCoverage package from deepTools (version 3.5.4) normalizing by counts per million (CPM). Count tables for genes were also generated from bam files using the featureCounts package from Subread (version 2.0.6) and differential gene expression analysis was performed using DESeq2 (version 1.40.2). Results were visualized using ggplot2 (version 3.4.4).

### AML cell line bulk ATAC-seq analysis

FASTQ files were demultiplexed with bcl2fastq and aligned to the hg38 reference genome using bowtie2 (version 2.5.2). Sam files were sorted, indexed, and converted to bam files with Samtools (version 1.18). For data visualization in a genome browser, bam files were converted to bigwig files using the bamCoverage package from deepTools (version 3.5.4) normalizing by counts per million (CPM). Peak calling was performed using MACS2 (version 2.2.9.1) with the flags --shift -100 and --extsize 200 to generate narrowPeak files. NarrowPeak files from DMSO and dTAG^V^-1-treated cells were then merged and converted to SAF format to generate a consensus peak set for each cell line. This consensus peak file and corresponding bam files were then processed using the featureCounts package from Subread (version 2.0.6) to generate a count table for ATAC peaks. Differential peak accessibility analysis was performed using DESeq2 (version 1.40.2). Results were visualized using ggplot2 (version 3.4.4). Transcription factor binding motif enrichment analysis was performed using Analysis of Motif Enrichment (AME) from The MEME Suite (version 5.5.0) to assess enrichment of motifs from the JASPAR CORE (2022) Vertebrates Non-Redundant database^73^.

### PRO-seq analysis

All custom scripts described herein are available on the AdelmanLab GitHub (https://github.com/AdelmanLab/NIH_scripts). Using a custom script (trim_and_filter_PE.pl), FASTQ read pairs were trimmed to 41bp per mate, and read pairs with a minimum average base quality score of 20 were retained. Read pairs were further trimmed using cutadapt (version 4.1) to remove adapter sequences and low-quality 3’ bases (–match-read-wildcards -m 20 -q 10). R1 reads, corresponding to RNA 3’ ends, were then aligned to the spiked in Drosophila genome index (dm6) using BWA, with those reads not mapping to the spike genome serving as input to the primary genome alignment step. Reads mapping to the hg38 reference genome were then sorted, via samtools (version 1.3.1 -n), and subsequently converted to bam files. The bam files are converted to bigwig files by bamCoverage of deepTools (version 3.5). For metagene plots, bigwig files of three replicates of each group and combined and averaged using WiggleTools.

### ChIP-seq analysis

For HA and H3K27ac ChIP-seq experiments, FASTQ files were demultiplexed with bcl2fastq and aligned to the hg38 reference genome using bowtie2 (version 2.5.2). Sam files were sorted, indexed and converted to bam files with Samtools (version 1.18). For data visualization in a genome browser, bam files were converted to bigwig files using the bamCoverage package from deepTools (version 3.5.4) normalizing by counts per million (CPM). Peak calling was performed using MACS2 (version 2.2.9.1). NarrowPeak files from DMSO and dTAG^V^-1-treated cells were then merged and converted to SAF format to generate a consensus peak set for each cell line. This consensus peak file and corresponding bam files were then processed using the featureCounts package from Subread (version 2.0.6) to generate a count table for ChIP peaks. CtBP2 ChIP-seq summary data (Bigwig files aligned to hg19) were downloaded from GSE236010 and converted to Bigwig files aligned to hg38 using the liftOver tool from UCSC. Bigwig files were analyzed with the computeMatrix and plotHeatmap packages from deepTools to assess the deposition of H3K27ac signal and association of CtBP2 at MECOM-regulated cisREs.

### Identification of MECOM-regulated *cis*-regulatory elements

Using external parental MUTZ-3 MECOM ChIP-seq data^30^, we performed peak calling using MACS2 as described above. We intersected the genomic coordinates of the ChIP-seq peaks with our ATAC-seq peak calls to identify MECOM-bound sites with open chromatin. To enrich for a set of sites with both strong MECOM-binding and a change in chromatin accessibility following MECOM degradation we filtered our overlapping sites using the following parameters: ATAC-seq (6hr dTAG^V^-1 vs DMSO) DESeq2 p-value <0.01 and MECOM ChIP-seq MACS2 peak Pscore>50 (PScore=-log10pvalue*10). This analysis resulted in 837 genomic intervals (MECOM cisRE network).

### Gene set enrichment analysis

We used GSEApy^74^ (https://github.com/zqfang/GSEApy) for all GSEA analyses to determine the enrichment of genes and cisREs under direct regulation of MECOM as determined by our MUTZ-3 dTAG studies in other *MECOM-FKBP12*^*F36V*^ cell line models. Significant enrichment of the gene and cisRE sets was determined using GSEAPrerank in which the RNA-seq and ATAC-seq data from UCSD-AML1, HNT-34, and OCI-AML4 cells treated with dTAG^V^-1 vs. DMSO were pre-ranked by log_2_ fold change. For enrichment analyses of MECOM-regulated cisREs, the genomic coordinates of ATAC-seq peaks were used in place of gene name. ATAC-seq peaks from all datasets were overlapped with the MUTZ-3 consensus cisRE network using the intersect package from bedtools (version 2.31.0) and renamed to the same genomic coordinates of the corresponding MUTZ-3 peak. GSEA was performed using 1,000 permutations to determine significance.

### Genome Regions Enrichment of Annotations Tool (GREAT) analysis

Differentially accessible ATAC-peaks that also overlapped with MECOM ChIP-seq peaks (837 site cisRE network) (**Fig. 2E**) were linked to genes based on proximity using Genome Regions Enrichment of Annotations Tool (GREAT; version 4.0.4). Each cisRE was associated to genomic loci using the “basal plus extension” mode with the following parameters: proximal 5kB upstream, 1kb downstream, plus distal up to 1000kb.

### Mass spectrometry data analysis

Mass spectrometry data were analyzed using the open-source Comet algorithm (release_2019010), following a previously established pipeline and a customized FASTA-formatted database^75–78^. This database included common contaminants and reversed sequences (Uniprot Human, 2021). The search parameters were set as follows: 50 PPM precursor tolerance, fully tryptic peptides, 0.02 Da fragment ion tolerance, and static modifications of TMTpro18 (+304.2071 Da) on lysine residues and peptide N-termini, as well as carbamidomethylation of cysteine residues (+57.0214 Da). Oxidation of methionine residues (+15.9949 Da) was included as a variable modification.

Peptide spectral matches were filtered to maintain a false discovery rate (FDR) of <1% using linear discriminant analysis with a target-decoy strategy. Further filtering ensured a protein-level FDR of 1% across the dataset, and proteins were grouped accordingly. Reporter ion intensities were corrected for TMT reagent impurities following the manufacturer’s specifications. MS2 spectra required a total signal-to-noise (S/N) sum of at least 180 across all reporter ions for quantification. For proteins, S/N measurements of corresponding peptides were summed and normalized to ensure consistent loading across all channels. Finally, protein abundance measurements were scaled such that the total summed S/N for each protein across all channels was set to 100, providing relative abundance measurements.

### CRISPR screen analysis

Quality control and enrichment analysis of CRISPR screen sequencing data was performed using MAGeCKFlute pipeline^79^ (version 0.5.9.5). Briefly, FASTQ files were mapped using the count function with the control norm-method. Enrichment of cisRE-targeting sgRNAs was calculated using the test function comparing either the sorted CD34^+^ population (CRISPRi screen) or the sorted CD34-low population (CRISPRa screen) to the plasmid DNA library, with the MAGeCK Robust Rank Algorithm (RRA) using non-targeting and AAVS1-targeting sgRNAs as negative controls.

### Single cell RNA-seq analysis

Filtered count matrices were downloaded from GEO (GSE235063) and analyzed according to Lambo et al. 2023^34^. Briefly, data were log normalized to 10,000 counts and scaled using Seurat^80^. Dimensionality reduction was performed by Uniform Manifold Approximation and Projection (UMAP) using the uwot package (version 0.1.1) after initialization using principal component analysis (PCA). Calculation of nearest neighbors (nn) was performed by applying the annoy algorithm. Inferred cell type, malignancy and other metadata were carried over from Lambo et al 2023^34^.

Malignant cells were selected according to their annotation. Signature scores were calculated using the Addmodulescore function in Seurat^80^ using published signatures of HSC and monocyte populations from Lambo et al. 2023^34^ and MECOM-regulated genes from this study (**Table S7**). Comparison of signature scores between MECOM positive and MECOM negative samples was performed using a Wilcoxon signed rank test adjusted for multiple testing correction using Benjamini Hochberg (BH) correction. Differential expression was calculated using MAST^81^ (version 1.16) as implemented within the findMarker feature in Seurat. Comparisons were performed by randomly taking the average over ten iterations of 1,000 randomly sampled cells from both samples expressing MECOM and samples not expressing MECOM to avoid uninformative p-values close to zero.

### Primary AML bulk RNA-seq analysis

Data were downloaded and processed as described in Lambo et al. 202334 Samples were deemed MECOM positive if expression was over 32 transcripts per million. Gene set enrichment was calculated using gene set variation analysis (GSVA v1.38.2)82 using an HSC signature derived from Lambo at al. 2023.

### Single cell ATAC-seq analysis

Filtered fragment files were downloaded from GEO (GSE235308) and remission samples were analyzed according to Lambo et al. 2023^34^. Briefly, cells were clustered using iterative latent semantic indexing (LSI)^83^ and annotations, including cell type labels, malignancy status, peak calling, and linked scRNA cells were transferred from the original publication. Linkage between scRNA profiles and scATAC profiles was also based on this metadata and were originally identified using Seurat findTransferAnchor^80^. Markov Affinity-based Graph Imputation of Cells (MAGIC; version 2.0)^84^ was used to impute weights based on identified nearest neighbors in the dimensionality reduction. Motif analysis was performed using Chromvar^85^ using annotations derived from cisBP^86^.

Subsequently, MECOM cisRE insertion scores were calculated by summing up insertions within peaks identified six hours post dTAG^V^-1 treatment (**Table S6**) and normalized by the total insertions in promoter regions. This was performed to correct for sequencing depth and differences in signal to noise ratio within cells. Lineages scores were defined using lineage defining peaks, which were derived from Lambo et al 2023^34^. Lineage scores were calculated for each cell separately by combining the total insertions within lineage defining peaks of each separate lineage (Myeloid, Lymphoid, Erythroid) and dividing this number by the total insertions of lineage defining peaks from the other lineages. Correlations between scores were calculated using Spearman correlation.

Trajectory analysis was performed using Monocle (version 3.0)^39^ between clusters defined by Seurat findClusters^80^. Trajectories were drawn between the cluster with the highest number of identified HSCs^80^, and CD34 positive cells and the cluster with the highest number of monocytes and CD14 positive cells. Cells along the cluster were binned in 100 bins of equal size and profiles were aggregated, z-score normalized and smoothened using a rolling mean across the trajectory. A heatmap of cluster scores was calculated using the mean score of each cluster and scaled using z-score normalization.

### Quantification and statistical analysis

Statistical tests and statistical significance are indicated in the figure legends. All error bars represent standard error of the mean unless otherwise indicated.

**Figure S1:**
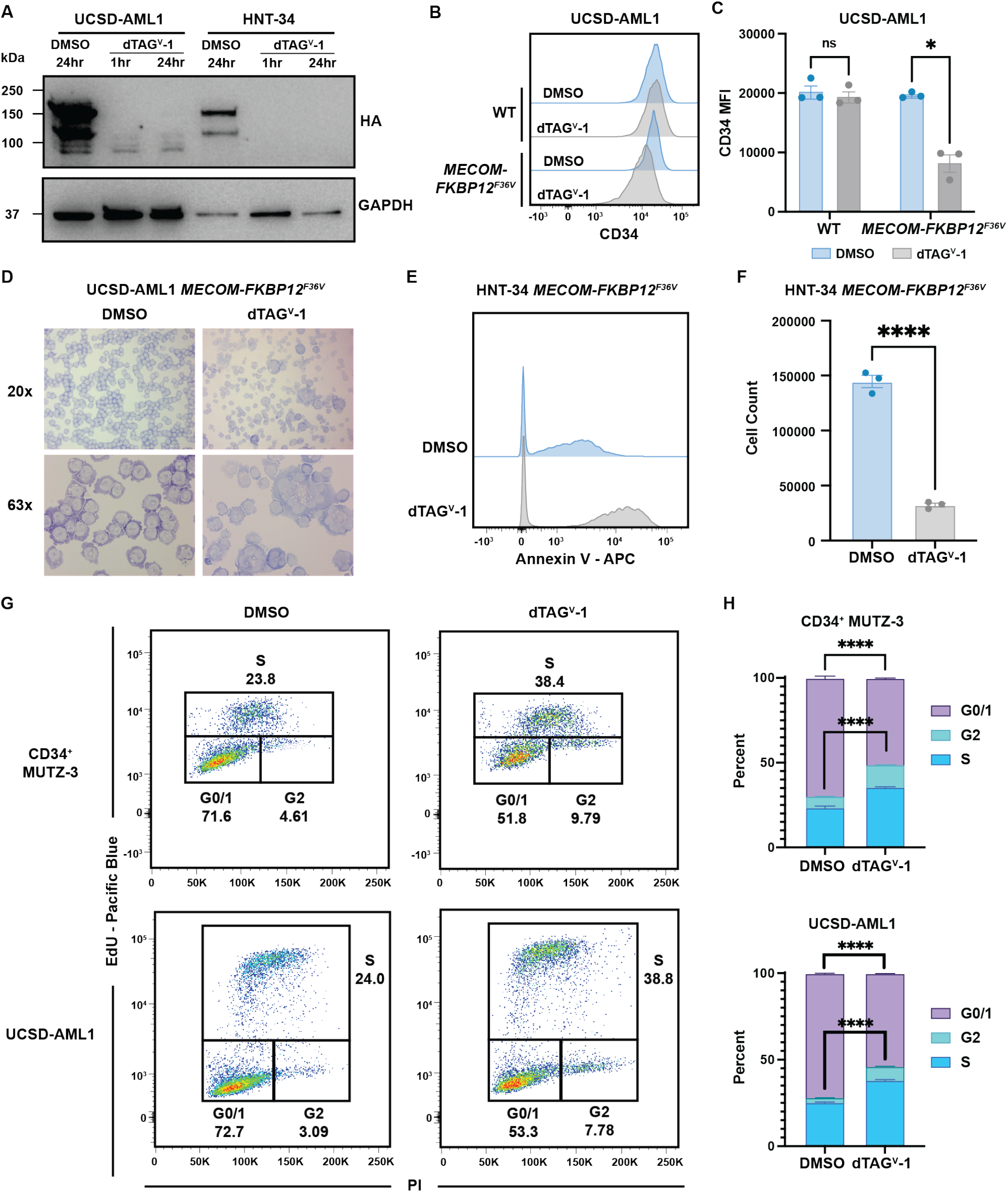
FKBP12^F36V^ degron enables targeted MECOM degradation in additional AML cell lines. **(A)** Time course western blot analysis of MECOM protein levels in UCSD-AML1 MECOM-FKBP12^F36V^ and HNT-34 MECOM-FKBP12^F36V^ cells treated with 500nM dTAG^V^-1 vs. DMSO. **(B-C)** Histogram of CD34 expression in UCSD-AML1 MECOM-FKBP12^F36V^ and WT UCSD-AML1 cells 6 days after treatment with 500nM dTAG^V^-1 vs. DMSO. n = 3 independent replicates, mean and SEM are shown. Two-sided Student t test was used for comparison. *p < 0.05, ns, not significant. **(D)** Confocal microscopy images of UCSD-AML1 MECOM-FKBP12^F36V^ cells 9 days after treatment with 500nM dTAG^V^-1 vs. DMSO following cytospin and May-Grünwald Giemsa staining. **(E)** Histogram showing Annexin-V staining of HNT-34 MECOM-FKBP12^F36V^ cells 3 days after treatment with 500nM dTAG^V^-1 vs. DMSO. n = 3 independent replicates. **(F)** Viable cell count by trypan blue exclusion of HNT-34 MECOM-FKBP12^F36V^ 6 days after treatment with 500nM dTAG^V^-1 vs. DMSO. n = 3 independent replicates, mean and SEM are shown. Two-sided Student t test was used for comparison. ****p < 0.0001. **(G-H)** Cell cycle analysis of CD34+ MUTZ-3 MECOM-FKBP12^F36V^ and UCSD-AML1 MECOM-FKBP12^F36V^ cells after treatment with 500nM dTAG^V^-1 vs. DMSO. **(G)** 48 hours post-treatment, cells were incubated with 10uM EdU for 2 hours then processed for flow cytometry analysis with PI staining. **(H)** Stacked bar plot comparing differences in cell cycle populations (G0/1, G2, S) of dTAG^V^-1 vs. DMSO treated samples. n = 3 independent replicates, mean and SEM are shown. Two-sided Student t test was used for comparison. ****p < 0.0001.

**Figure S2:**
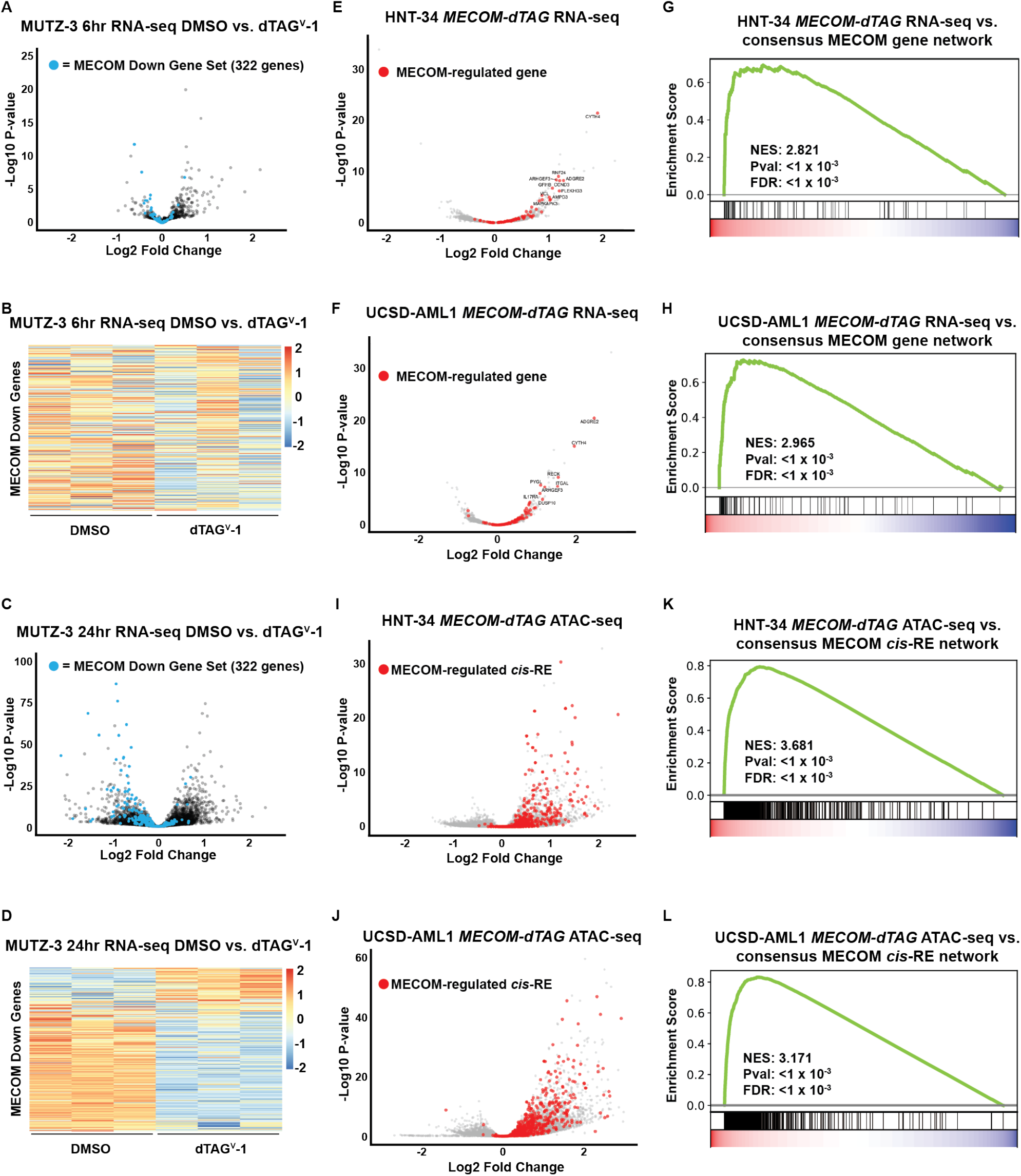
MECOM-regulated gene and chromatin networks are conserved in UCSD-AML1 and HNT-34 cell line models. **(A-B)** Volcano plot and heatmap representing changes in gene expression assessed via bulk RNA-seq of MUTZ-3 dTAG cells treated with dTAG^V^-1 or DMSO for 6 hours (n=3). MECOM down genes^14^ are highlighted in blue data points. **(C-D)** Volcano plot and heatmap representing changes in gene expression assessed via bulk RNA-seq of MUTZ-3 dTAG cells treated with dTAG^V^-1 or DMSO for 24 hours (n=3). MECOM down genes are highlighted in blue data points. **(E-H)** Volcano plots showing changes in gene expression assessed via bulk RNA-seq of UCSD-AML1 MECOM-FKBP12^F36V^ and HNT-34 MECOM-FKBP12^F36V^ cells treated with dTAG^V^-1 or DMSO for 6 hours (n=3) and gene set enrichment analysis (GSEA) compared to MECOM network genes. MECOM network genes as identified from MUTZ-3 experiments in **Fig. 2** are highlighted in red data points. **(I-L)** Volcano plots showing changes in chromatin accessibility assessed via ATAC-seq of UCSD-AML1 MECOM-FKBP12^F36V^ and HNT-34 MECOM-FKBP12^F36V^ cells treated with dTAG^V^-1 or DMSO for 6 hours (n=3) and gene set enrichment analysis (GSEA) of ATAC-peaks compared to MECOM network cisREs. MECOM network cisREs as identified from MUTZ-3 experiments in **Fig. 2** are highlighted in red data points.

**Figure S3:**
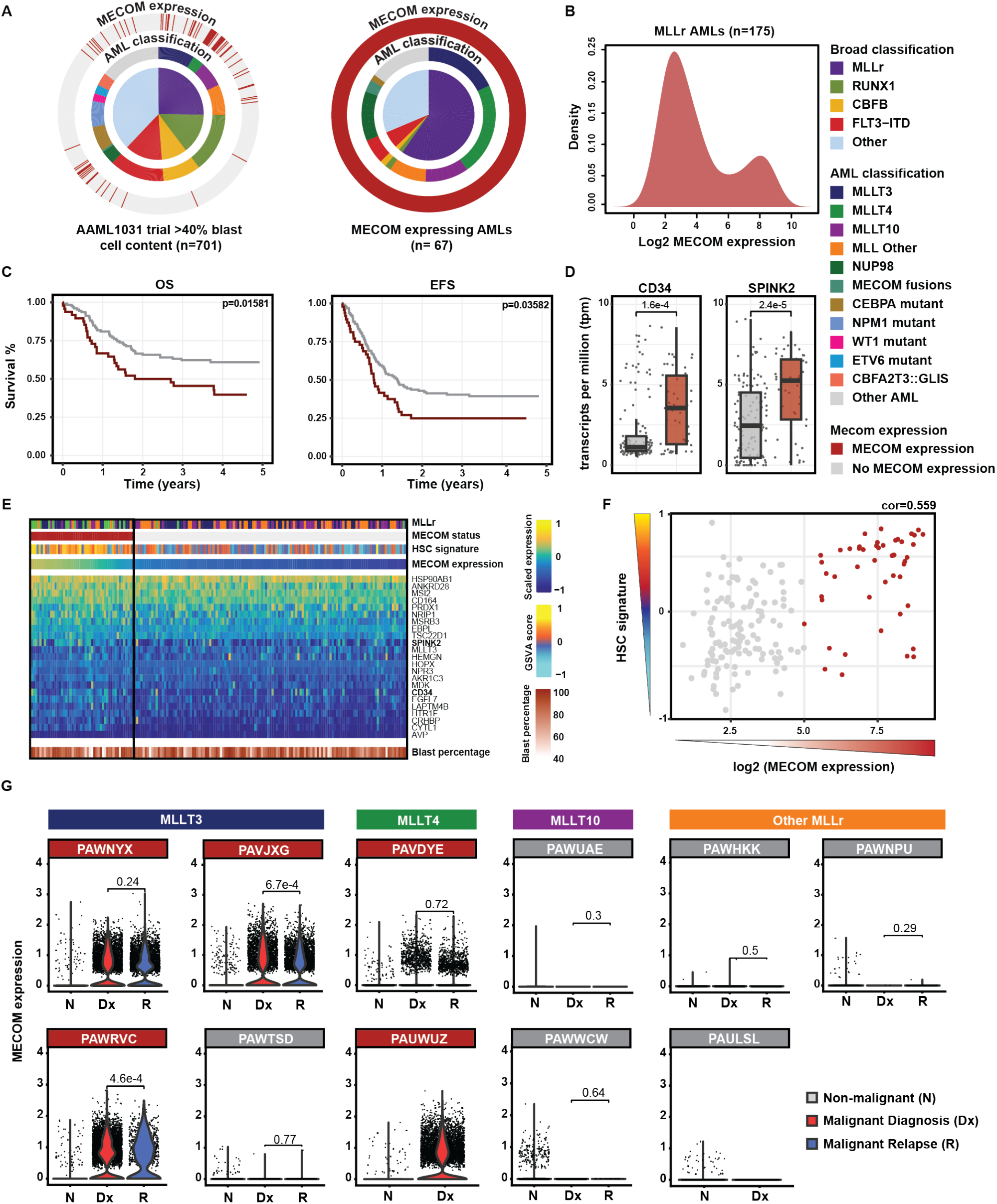
Primary MLL-rearranged AML cohort is stratified by MECOM expression status. **(A)** Representation of patient samples at diagnosis sequenced using bulk RNA-seq as part of the AAML1031 trial (n=701) (Aplenc et al 2020^33^, Lambo et al 2023^34^). (Left) From inside to outside, the inner circle shows a broad classification used in Lambo et al 2023^34^, the second ring shows a more detailed cytogenetic classification of the MLLr subgroup and other AMLs and the outer ring shows whether MECOM was found to be expressed (log_2_ expression > 5). (Right) Graph showing the same classification of all samples having MECOM expression. Only samples with a blast cell content over 40% were included. **(B)** Expression of MECOM across the MLLr leukemias included in the AAML1031 cohort. x-axis represents log_2_ transformed expression value, y-axis represents the kernel density. **(C)** Kaplan meier survival curves showing five-year OS and five-year EFS split by MLLr leukemias expressing MECOM (n=49) and MLLr leukemias not expressing MECOM (n=126). **(D)** Bar chart showing the expression of HSC-associated genes CD34 and SPINK2 in samples stratified by MECOM expression. BH-adjusted p-values were calculated using two-sided Wilcoxon signed-rank tests. **(E)** Heatmap showing the expression pattern of HSC signature genes derived from normal HSCs (Lambo et al 2023^34^) across 175 MLLr samples within the AAML1031 cohort. Signature scores were calculated using GSVA. **(F)** Scatter plot showing the correlation between HSC signature scores derived using GSVA and the log_2_ expression of MECOM. **(G)** Violin plots showing the expression of MECOM across individual cells of 11 MLLr samples included in the scRNA cohort of the AAML1031 trial. Cells were divided by non-malignant as described in (Lambo et al 2023^34^), samples taken at diagnosis and at relapse. BH adjusted p-values were calculated using two-sided Wilcoxon signed-rank tests.

**Figure S4:**
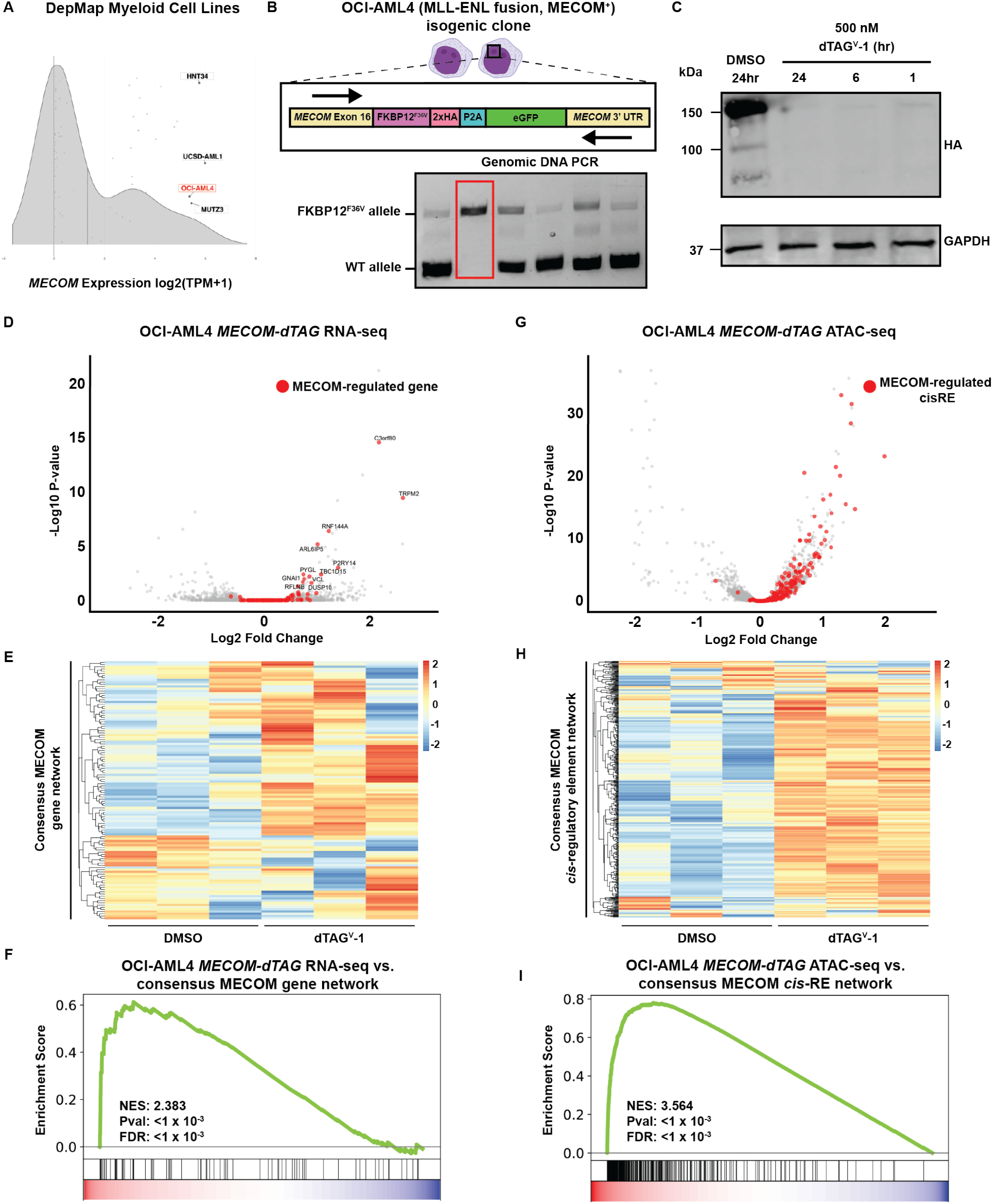
MECOM-regulated gene and chromatin networks are conserved in MLL-rearranged, MECOM+ OCI-AML4 cells. **(A)** Depiction of myeloid cell lines from The Cancer Dependency Map and their relative MECOM expression. **(B)** Genomic DNA PCR strategy to screen for MECOM-FKBP12^F36V^ biallelically-tagged OCI-AML4 isogenic clones. OCI-AML4 cells lack an activating translocation or rearrangement at the *MECOM* locus, thus requiring both *MECOM* alleles to be tagged with an FKBP12^F36V^ degron. PCR primers flanking the C-terminus of *MECOM* were used to identify a biallelically-tagged, isogenic clone (outlined in red). **(C)** Time course western blot analysis of MECOM protein levels in OCI-AML4 MECOM-FKBP12^F36V^ clone treated with 500nM dTAG^V^-1 vs. DMSO. **(D)** Volcano plot representing changes in gene expression assessed via RNA-seq of OCI-AML4 MECOM-FKBP12^F36V^ clone treated with dTAG^V^-1 vs. DMSO for 6 hours (n=3). MECOM network genes as identified from MUTZ-3 experiments in **Fig. 2** are highlighted in red data points. **(E)** Heatmaps displaying differential expression of individual MECOM network genes in DMSO and dTAG^V^-1 conditions from experiments in **Fig. S4D. (F)** Gene set enrichment analysis (GSEA) of differentially expressed genes from experiments in **Fig. S4D** compared to MECOM network genes. **(G)** Volcano plot representing changes in chromatin accessibility assessed via ATAC-seq of OCI-AML4 MECOM-FKBP12^F36V^ clone treated with dTAG^V^-1 vs. DMSO for 6 hours (n=3). MECOM-regulated *cis*-regulatory elements as identified from MUTZ-3 experiments in **Fig. 2** are highlighted in red. **(H)** Heatmap displaying differential accessibility of individual MECOM-regulated *cis*-regulatory elements in DMSO vs. dTAG^V^-1 samples from experiments in **Fig. S4G. (I)** Gene set enrichment analysis (GSEA) of differentially accessible ATAC-peaks from experiments in **Fig. S4G** compared to MECOM-regulated *cis*-regulatory elements.

**Figure S5:**
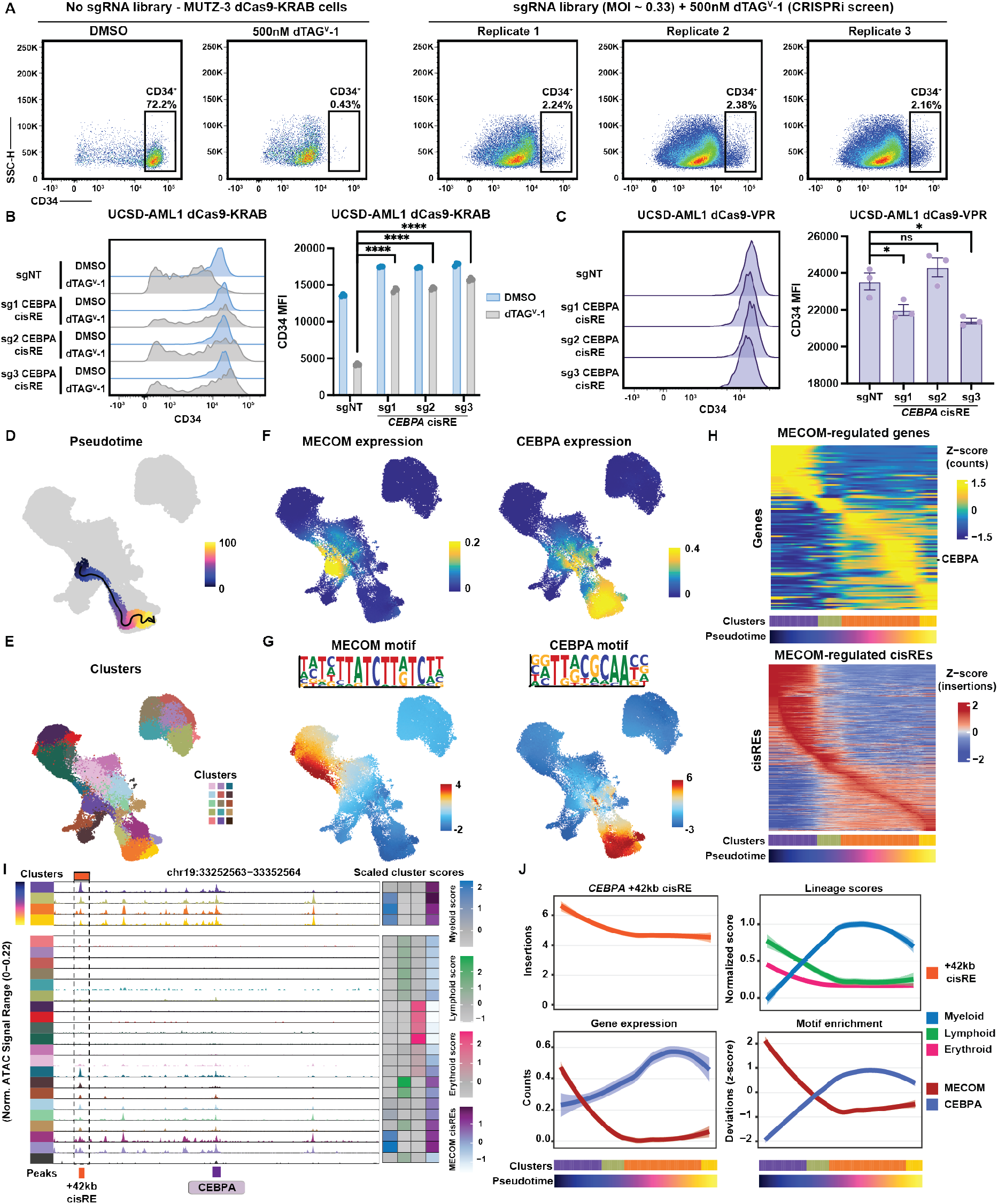
CEBPA cisRE function is conserved in other AML cells. **(A)** Flow cytometry plot from MUTZ-3 CRISPRi screen 14 days in culture. The no sgRNA library condition compared to sgRNA library transduced condition demonstrates the relative enrichment of the phenotypically rescued, CD34+ cells in sgRNA library expressing cells. **(B)** Orthogonal validation of CRISPRi screen in UCSD-AML1 cells. UCSD-AML1 MECOM-FKBP12^F36V^ dCas9-KRAB cells were infected with sgRNA-expressing lentiviruses targeting either the CEBPA cisRE or a non-targeting (NT) sequence. 48 hours after transduction, cells were treated with 500nM dTAG^V^-1 vs. DMSO. (Left) Histogram shows CD34 expression at day 9. (Right) Percentage of CD34+ cells at day 9. n = 3 independent replicates, mean and SEM are shown. Two-sided Student t test was used for comparison. ****p < 0.0001. **(C)** Orthogonal validation of CRISPRa screen in UCSD-AML1 cells. UCSD-AML1 dCas9-VPR cells were infected with sgRNA-expressing lentiviruses targeting either the CEBPA cisRE or a non-targeting (NT) sequence. (Left) Histogram shows CD34 expression at day 9. (Right) Percentage of CD34+ cells at day 9. n = 3 independent replicates, mean and SEM are shown. Two-sided Student t test was used for comparison. *p < 0.05, ns, not significant. **(D)** UMAP showing a trajectory inferred using Monocle from inferred HSCs to inferred Monocytes (see also **Figure 4**). **(E)** Seurat identified clusters in the scATAC-seq data. Each color represents a different cluster. **(F)** Expression of *MECOM* and *CEBPA* from scaled counts derived from linked scRNA-seq data across cells from remissions. **(G)** UMAP showing the scaled TF motif enrichment scores (ChromVAR deviation scores) of MECOM and CEBPA motifs across all remission cells. **(H)** Heatmaps showing the scaled expression of identified genes (n=122) and cisREs (n=837) directly regulated by MECOM **(Figure 2)** along the pseudotime shown in **(E)**. Each column represents an aggregated minibulk from cells across the inferred pseudotime (100 bins total). Gene expression scores are counts derived from linked scRNA samples, peak insertions were normalized by TSS insertions and Tn5 bias. Both gene expression and chromatin accessibility were scaled across all cells in the pseudotime. **(I)** Normalized ATAC signal across clusters identified in **(B)** within 100 kB around the *CEBPA* locus (chr19:33252563-33352564). The *CEBPA* +42kb cisRE is shown in orange. The *CEBPA* promoter is shown in purple. The four clusters taken for the pseudotime analysis were plotted separately on top and other clusters were sorted according to lineage. The heatmap on the right shows the aggregated scores for each lineage and MECOM-regulated cisRE insertions (identified in **Figure 4B-C**). **(J)** Plots showing the normalized insertions in the *CEBPA* +42kb cisRE (top left), lineage scores (top right), MECOM and CEBPA expression (bottom left) and MECOM and CEBPA TF motif enrichment (bottom right) across the pseudotime identified in **(G)** and **Figure 4**. Insertions, signature scores, expression and TF motif enrichment were scaled across all cells in the pseudotime and smoothened using LOESS.

**Figure S6:**
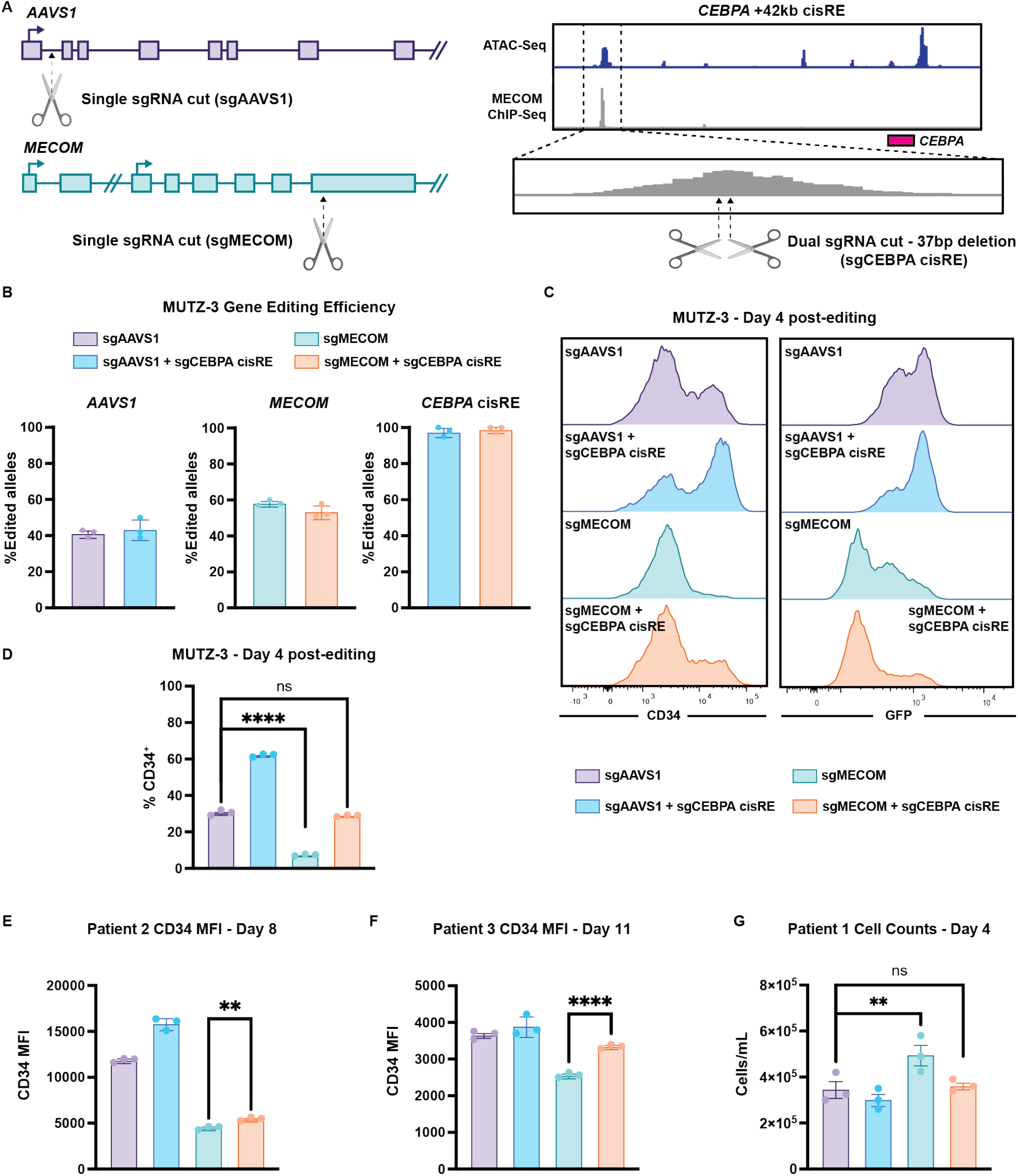
Proof-of-concept gene editing strategy to inactivate the CEBPA cisRE with Cas9 nuclease. **(A)** Schematic showing gene editing strategy to inactivate *AAVS1, MECOM*, and *CEBPA* cisRE. (Left) *AAVS1* and *MECOM* were targeted with single sgRNAs targeting the early coding sequence (CDS). (Right) The *CEBPA* cisRE was targeted with two sgRNAs proximal to the summit of the MECOM ChIP-seq peak to create a 37bp inactivating deletion. In a proof-of-concept experiment for this gene editing approach, WT MUTZ-3 cells were electroporated with Cas9 RNPs targeting *AAVS1* and *MECOM* with and without *CEBPA* cisRE targeting. **(B)** Efficiency of gene editing in MUTZ-3 cells at the *AAVS1, MECOM*, and *CEBPA* (cisRE) loci. Editing estimated using Sanger sequencing of amplicons followed by sequence trace decomposition analysis with the ICE tool^57^. For *CEBPA* cisRE, only deletions resulting from dual guide cleavage were counted. n = 3 independent replicates, mean and SEM are shown. **(C-D)** Immunophenotypic analysis of MUTZ-3 cells 4 days post-electroporation with Cas9 RNPs. **(C)** Histogram showing CD34 expression assessed by flow cytometry. **(D)** CD34 expression measured by MFI. n = 3 independent replicates, mean and SEM are shown. Two-sided Student t test was used for comparison. ****p < 0.0001. ns, not significant. **(E-F)** CD34 expression measured by MFI from experiments on primary leukemia patient samples (patients 2-3, **Table S9**) in **Fig. 6**.n = 3 independent replicates, mean and SEM are shown. Two-sided Student t test was used for comparison. **p < 0.01, ****p < 0.0001. **(G)** Viable cell counts by trypan blue exclusion in primary leukemia sample (patient 1, **Table S9**) 8 days post-electroporation from **Fig. 6**. n = 3 independent replicates, mean and SEM are shown. Two-sided Student t test was used for comparison. **p < 0.01, ns, not significant.

**Figure S7:**
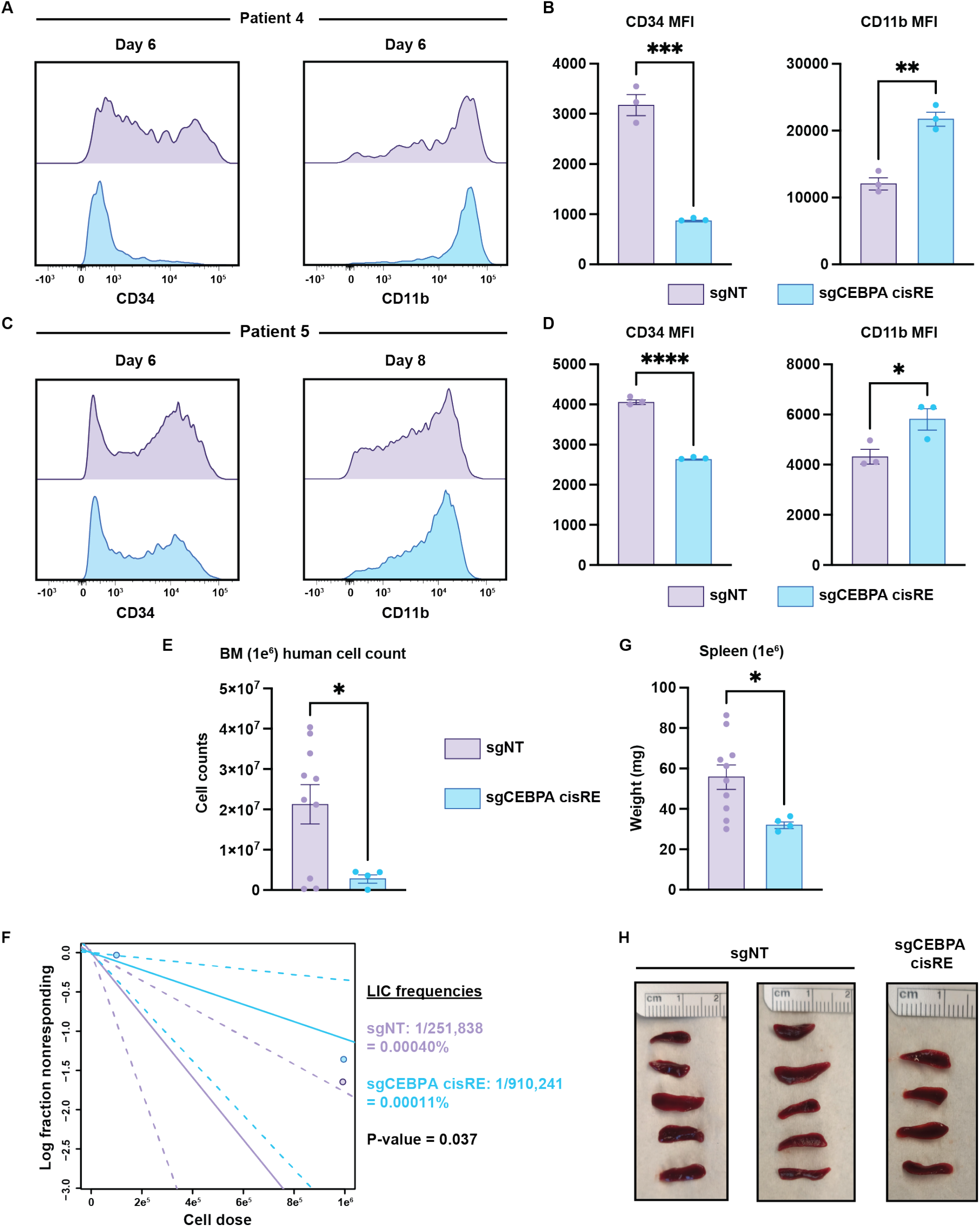
Activation of CEBPA cisRE induces myeloid differentiation of additional primary AML samples. **(A-B)** Immunophenotypic analysis of primary leukemia sample (patient 4, **Table S9**) 6 days post-electroporation with dCas9-VPR mRNA and sgRNAs targeting *CEBPA* cisRE or NT **(A)** Histograms showing CD34 and CD11b expression assessed by flow cytometry. **(B)** CD34 and CD11b expression measured by MFI. n = 3 independent replicates, mean and SEM are shown. Two-sided Student t test was used for comparison. **p < 0.01, ***p < 0.001. **(C-D)** Immunophenotypic analysis of primary leukemia sample (patient 5, **Table S9**) 6-8 days post-electroporation with dCas9-VPR mRNA and sgRNAs targeting *CEBPA* cisRE or NT **(C)** Histograms showing CD34 (day 6) and CD11b (day 8) expression assessed by flow cytometry. **(D)** CD34 and CD11b expression measured by MFI. n = 3 independent replicates, mean and SEM are shown. Two-sided Student t test was used for comparison. ****p < 0.0001, *p < 0.05. **(E)** Quantification of human cells in the bone marrow of mice transplanted with 1e^6^ primary AML cells. Viable cell counts were determined by trypan blue exclusion. Cell quantities were calculated based on total cell counts observed in the bone marrow and the corresponding human cell chimerism (hCD45+). n = 4-10 xenotransplant recipients as shown, mean and SEM are shown. Two-sided Student t test was used for comparison. *p < 0.05. **(F)** Quantification of leukemia initiating cell (LIC) frequencies based on bone marrow engraftment rates using the ELDA software^51^. Engraftment of leukemia cells was considered successful if human cell chimerism in the bone marrow was >5%. **(G-H)** Weight (mg) and images of spleens from mice transplanted with 1e^6^ primary AML cells. n = 4-10 xenotransplant recipients as shown, mean and SEM are shown. Two-sided Student t test was used for comparison. *p < 0.05.

